# Structural variants are enriched in deleterious visible phenotypes in *Drosophila*

**DOI:** 10.1101/2025.08.15.670616

**Authors:** Alejandra Samano, Matthew Musat, Mihir Junaghare, Asad Ahmad, Mehlum Ali, Sebastian Alves, Sreeram Pasupuleti, Jelisha Perera, Omar Saada, Brady Sabido, Trevor Smith, Sophie Walz, Mahul Chakraborty

**Affiliations:** Department of Biology, Texas A&M University, College Station, TX 77843

## Abstract

Genome structural variants (SVs) comprise a sizable portion of functionally important genetic variation in all organisms; yet, many SVs evade discovery using short reads. While long-read sequencing can find the hidden SVs, the role of SVs in variation in organismal traits remains largely unclear. To address this gap, we investigate the molecular basis of 50 classical phenotypes in 11 *Drosophila melanogaster* strains using highly contiguous *de novo* genome assemblies generated with Oxford Nanopore long reads. These assemblies enabled the creation of a pangenome graph containing comprehensive, nucleotide-resolution maps of SVs, including complex rearrangements such as the interchromosomal inverted duplication Dp(2;4)eyD and large tandem duplications at the *Bar* locus. We uncovered new candidate causal mutations for 15 phenotypes and new molecular alleles for 2 mutations comprising tandem duplications, transposable element (TE) insertions, and indels. For example, we mapped the tarsal joint defect *Ablp^eyD^* to an 8 kb Roo retrotransposon insertion into an intergenic enhancer, a finding validated via CRISPR-Cas9. The wing vein phenotype *plexus* (px^1^) was linked to a 1.5 kb partial tandem gene duplication, and the century-old *Curved* (c^1^) wing phenotype was linked to a 7.5 kb DM412 retrotransposon inserted into the coding sequence of the muscle protein gene *Strn-Mlck*. We also unveiled 8 SV alleles of previously identified causal genes, including previously uncharacterized SVs underlying the extensively studied white and yellow phenotypes. Overall, 67.4% of the genes causing phenotypic changes harbored candidate SVs over 100 bp, whereas only 28% is expected based on euchromatic SVs. Our data, based on the 50 *Drosophila* phenotypes, 44 of which are strongly deleterious, suggests a disproportionately larger contribution of SVs to deleterious changes in visible phenotypes in *Drosophila*.

## Introduction

Understanding the mutational basis of phenotypic differences between individuals or species is a fundamental puzzle in biology. Mutations that cause large or perceptible changes in phenotypes play an important role in adaptive evolution, agriculture, and medical genetics (Dittmar et al. 2016; Marian 2020). Genetic mapping approaches, empowered by advances in sequencing and genotyping methods, have enabled the discovery of several variants with large effects on phenotypes. However, these mapping studies often focus on small mutations such as single nucleotide polymorphisms (SNPs) and small indels in non-repetitive sequences. Genome structural variants (SVs) resulting from duplication, transposition, deletion, insertion, or inversion of sequences alter more nucleotides and are more likely to affect gene function than SNPs. Recent discoveries of previously hidden SVs associated with phenotypes by long-read sequencing further suggest that SVs may explain a portion of phenotypic variation unexplained by SNPs (Merker et al. 2018). SVs also tend to segregate at lower frequencies than SNPs, consistent with the idea that they are more often deleterious and subject to stronger purifying selection (Chakraborty et al. 2019; Abel et al. 2020; Collins et al. 2020; Collins and Talkowski 2025). These observations suggest SVs would account for larger phenotypic changes affecting fitness more often than small variants, including SNPs. However, the relative prevalence of SVs among the causal mutations for large changes in organismal traits remains unclear, obscuring the significance of SVs in diseases and adaptive evolution.

Visible phenotypic changes are a powerful model for uncovering genotype-phenotype relationships (Sax 1923; Koornneef et al. 1983; Long et al. 1995; Doebley 2004). Early geneticists leveraged visible phenotypes as markers to construct genetic maps and discover fundamental principles of genetics (Bateson et al. 1905; Sturtevant 1913). This included Mendel’s studies on pea plants (Mendel), discoveries from T.H. Morgan and his colleagues (Morgan 1910; Bridges 1922; Muller 1927), and Barbara McClintock’s work in maize (McCLINTOCK 1950). In particular, delineating mutations underlying morphological variation has helped us understand evolution within and between species as well as elucidate genetic mechanisms of pathological conditions (Wild et al. 1997; Jeong et al. 2008; Chan et al. 2010; Imsland et al. 2012; Ghodsinejad Kalahroudi et al. 2014; Van’t Hof et al. 2016). A systematic inquiry into the role of SVs and SNPs on the molecular basis for morphological changes in a species can help elucidate the molecular properties of mutations underlying such phenotypic changes. However, a study examining the role of SVs in a set of phenotypic changes is still lacking.

The model organism *Drosophila melanogaster* has an extensive collection of phenotype markers, many of which are deleterious to the organism. These classic mutant phenotypes were initially selected for genetics studies without knowing the molecular nature of the underlying mutation. Although transposable elements (TEs) are known to underlie several visible phenotypes (Green 1988; Sankaranarayanan 1988), the overall prevalence of SVs among *D. melanogaster* visible mutations remains unknown. Thus, examining the involvement of SNPs and SVs in these trait variations can provide insight into the relative role of SVs in phenotypic variation and help elucidate the biological basis of their deleterious fitness effects. We investigated the molecular basis of 50 visible phenotypic changes in 11 strains (Supplementary Table 1). In particular, we sequenced the 11 genomes using Oxford Nanopore long reads and assembled a high-quality genome for each strain. We employed a pangenomic approach, constructing a graph-based representation of variation across the 11 assemblies relative to the ISO1 reference, to create a comprehensive variant map and elucidate the molecular basis of variants associated with phenotypic changes.

## Results

### De novo genome assembly

We collected deep coverage (average coverage 90×, genome size or G = 140 Mb) ONT long-read sequences for 11 genomes carrying 50 visible mutations: 45 mutations are spontaneous, four are caused by X-ray irradiation, and one is due to a chemical mutagen (Supplementary Table 6). Although the average read accuracy of ONT long reads is 85-95%, the high read coverage (55-141×) used to assemble the genomes reported here provides high consensus accuracy for any genomic position (Sereika et al. 2022; Kolmogorov et al. 2023). A uniform coverage of mapped long reads across the assemblies suggests that our assemblies are free of large-scale assembly errors (Supplementary Fig. 1). All chromosome arms in our assemblies are represented by highly contiguous sequences (median assembly contig N50 = 23.4 Mbp), which include all of the euchromatin and a portion of pericentromeric heterochromatin in single contigs (Fig. 1a, Supplementary Fig. 2). However, the balancer chromosomes in two strains (1570 and 6027) are represented by fragmented assemblies due to the challenges of assembling heterozygous sequences with long error prone reads (Li and Durbin 2024). The number of Dipteran Benchmarking Universal Single Copy Orthologs (BUSCOs) in these genomes (complete BUSCO scores 98.6 – 99.5) and contiguity are comparable to the reference genome ISO1 (complete BUSCO score 98.8), further underscoring the high quality of the assemblies (Table 1).

**Figure 1.**
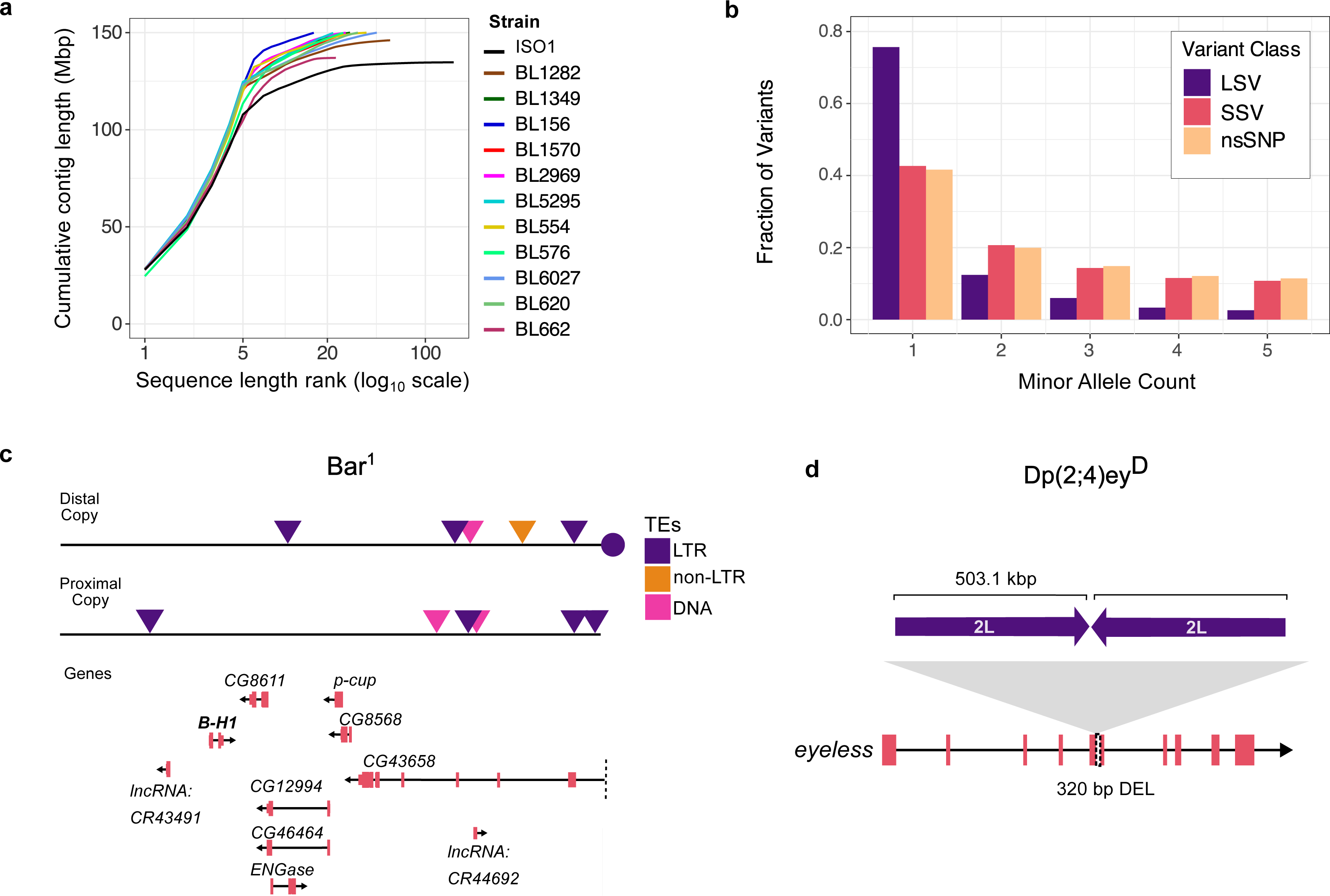
a. Contiguity plot comparison between the ISO1 contig-level assembly and our 11 genome assemblies. b. Minor allele counts for large SVs (LSVs; >100 bp), small SVs (SSVs; 10-100 bp), and nonsynonymous SNPs (nsSNPs). c. Gene and TE content of the duplicated sequences at the *Bar* locus (ISO1 coordinate X:17,334,493-17,537,993), which consists of nine complete genes and one truncated gene (*CG4368*). A *Roo* element (circle) separates the two copies, consistent with the hypothesized mechanism of TE-mediated duplication. d. The Dp(2;4)ey^D^ mutation is a translocation-duplication of a large sequence from chromosome 2L into an exon of the *eyeless* gene on chromosome 4, replacing 320 bp of coding sequence.

**Table 1.** Assembly Statistics.

|  | Contig Assembly |  |  | Scaffolded Assembly |  | Diptera BUSCOs |
| --- | --- | --- | --- | --- | --- | --- |
|  | Size (Mbp) | # contigs | Contig N50 (Mb) | # scaffs | Scaffold N50 (Mb) | Complete |
| BL1570 | 159.8 | 97 | 21.8 | 31 | 27.1 | 99.4 |
| BL6027 | 151.4 | 69 | 23.9 | 48 | 28.3 | 98.6 |
| BL662 | 135.7 | 22 | 21.8 | 13 | 26.6 | 99.2 |
| BL156 | 154.9 | 41 | 23.8 | 22 | 29.1 | 99.4 |
| BL1282 | 146 | 56 | 24.1 | 33 | 27.4 | 99.4 |
| BL2969 | 151.8 | 55 | 24 | 36 | 29.5 | 99.3 |
| BL554 | 150.4 | 51 | 24.1 | 27 | 28.6 | 99.4 |
| BL5295 | 154.3 | 54 | 24.3 | 31 | 27.1 | 99.4 |
| BL576 | 151.8 | 44 | 21.7 | 21 | 29 | 99.5 |
| BL620 | 151.9 | 51 | 24.3 | 28 | 28.9 | 99.4 |
| BL1349 | 153.4 | 78 | 24.2 | 55 | 29.3 | 99.4 |

### Landscape of genetic variation in 11 genomes

We identified mutations by comparing the genome assemblies of each strain to the ISO1 reference genome (Hoskins et al. 2015). To map genetic variation, we constructed a pangenome graph that captures all classes of mutations (Hickey et al. 2023). Unlike traditional approaches that compare each genome to a single linear reference, a pangenome graph represents sequences as nodes in a network, allowing us to identify both shared and unique variants, including SNPs, small indels, and SVs across all genomes (Eizenga et al. 2020; Sirén et al. 2021). While we also performed pairwise genome alignments and read mapping to the reference genome to validate genotypes at candidate loci, the pangenome graph provides a comprehensive and accurate view of genetic variation. Focusing on the euchromatic regions of the five major chromosome arms (2L, 2R, 3L, 3R, and X), we identified 53,337 small structural variants (SSVs; 10-100 bp) and 11,587 large structural variants (LSVs; >100 bp) across the 11 genomes. Of the LSVs, 7,156 were associated with TEs. We also identified 1.62 million SNPs, of which 5.2% affect coding exons.

We examined the minor allele frequency distribution of LSVs, SSVs, and nonsynonymous SNPs (nsSNPs). LSVs are significantly skewed towards lower frequencies than nsSNPs, a pattern likely driven by TEs (p-value < 2.2 × 10^-16^, 𝝌^2^ test between frequency distributions of LSVs and nsSNPs) (Fig. 1b) (Cridland et al. 2013; Chakraborty et al. 2019). Although SSVs also showed a skew toward lower frequencies, their distribution is more similar to that of nsSNPs. These patterns align with previous population genomics studies, which suggest that SVs are subject to stronger purifying selection, likely due to their more deleterious effects (Cridland et al. 2013; Chakraborty et al. 2019; Samano et al. 2025).

### Assembly of large, complex SVs

Large and repetitive SVs are often difficult to resolve at the molecular level (Treangen and Salzberg 2011). To assess the capacity of our assemblies to characterize such mutations, we analyzed two visible mutations known to be linked to large duplications. Strain 2969 carries the *Bar^1^* allele, an X-linked mutation that causes a slit-eye phenotype in males and homozygous females (Tice 1914). This phenotype was hypothesized to result from a tandem duplication caused by unequal crossing-over at a *Roo* element (Sturtevant 1925; Muller 1936), a hypothesis later supported by cloning and short-read sequencing of the *Bar* locus (Tsubota et al. 1989; Miller et al. 2016). Our genome assembly of strain 2969 captures both copies of the 203.5 kb duplicated region. The breakpoints match prior studies and include a *Roo* element between the two copies, supporting the TE-induced duplication model (Supplementary Fig. 3). The duplicated segment contains seven complete protein-coding genes, including *BarH1*, which is associated with the *Bar* eye phenotype (Kojima et al. 1993), as well as two long non-coding RNAs (lncRNAs), and one truncated gene (Fig. 1c). Notably, the two copies show substantial sequence divergence, including TE insertions unique to one copy (Fig. 1c). We also analyzed strain 662, which carries the *Dp(2;4)ey^D^* mutation, an X-ray induced translocation-duplication resulting in reduced or absent eyes (Hochman et al. 1964). Consistent with earlier cloning experiments (Kronhamn et al. 2002), we identified the 503.1 kb sequence from chromosome 2L, which was duplicated, with one copy inverted and inserted into the *eyeless* gene on the 4th chromosome (Fig. 1d, Supplementary Fig. 4). This translocation disrupts a coding exon and removes 320 bp of coding sequence from *ey*. Unlike the *Bar^1^*duplicates, the duplicated sequences of the *Dp(2;4)ey^D^* mutation show fewer sequence differences.

### Discovery of previously uncharacterized mutations

Strains carrying the *Bar^1^* and *Dp(2;4)ey^D^* mutations were included in this study due to their known association with large genomic rearrangements. In contrast, the other strains were selected without prior consideration of the molecular basis of their visible phenotypes. We first examined the molecular nature of mutations in genes previously linked to phenotypes, such as the *white* eye color mutation, as hidden SVs can mislead inferences of causal mutations for phenotypic changes (Chakraborty et al. 2019; Ebert et al. 2021; Fadaie et al. 2021)(Table 2). If the previously documented mutation was not present, we checked for the presence of another disruptive mutation in the same gene. Among the 50 phenotypes examined, we found that in 31 cases, the only candidate mutation present in the gene was the previously documented allele. We identified 19 previously uncharacterized mutations, including two found alongside the documented allele and one that contradicted the mutation type reported in FlyBase (Öztürk-Çolak et al. 2024). For 15 phenotypes with no prior molecular characterization, we used our pangenome graph to identify candidate mutations. However, four of these phenotypes were yet to be mapped on the *D. melanogaster* genome, so we combined our comprehensive variant map with additional genetic mapping experiments to identify the candidate mutations for the phenotypes.

**Table 2.**
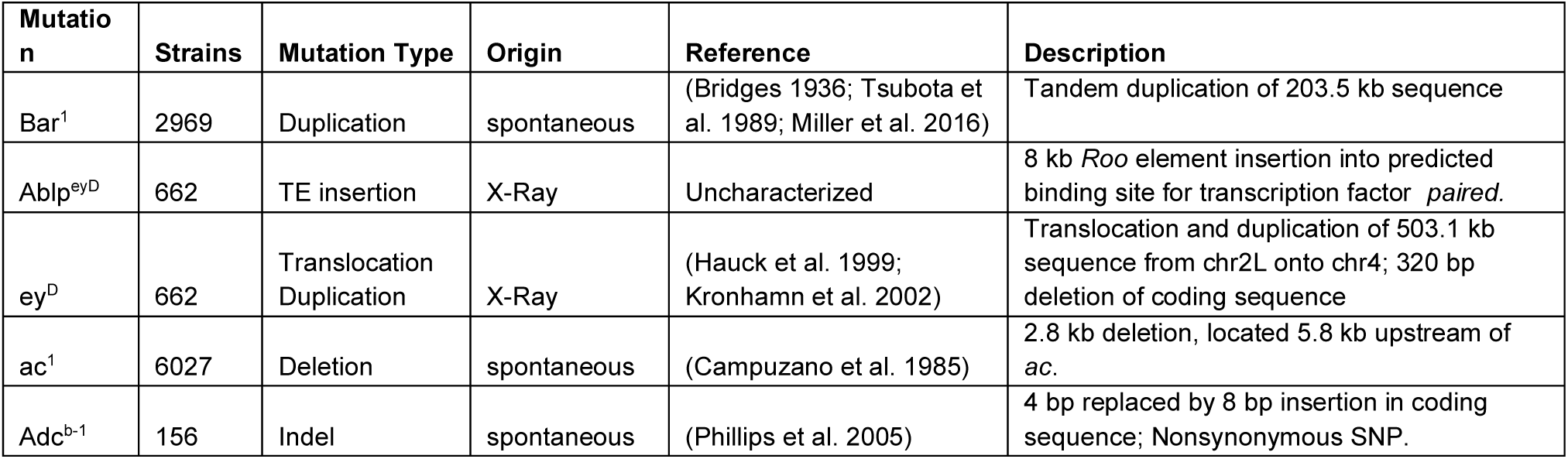
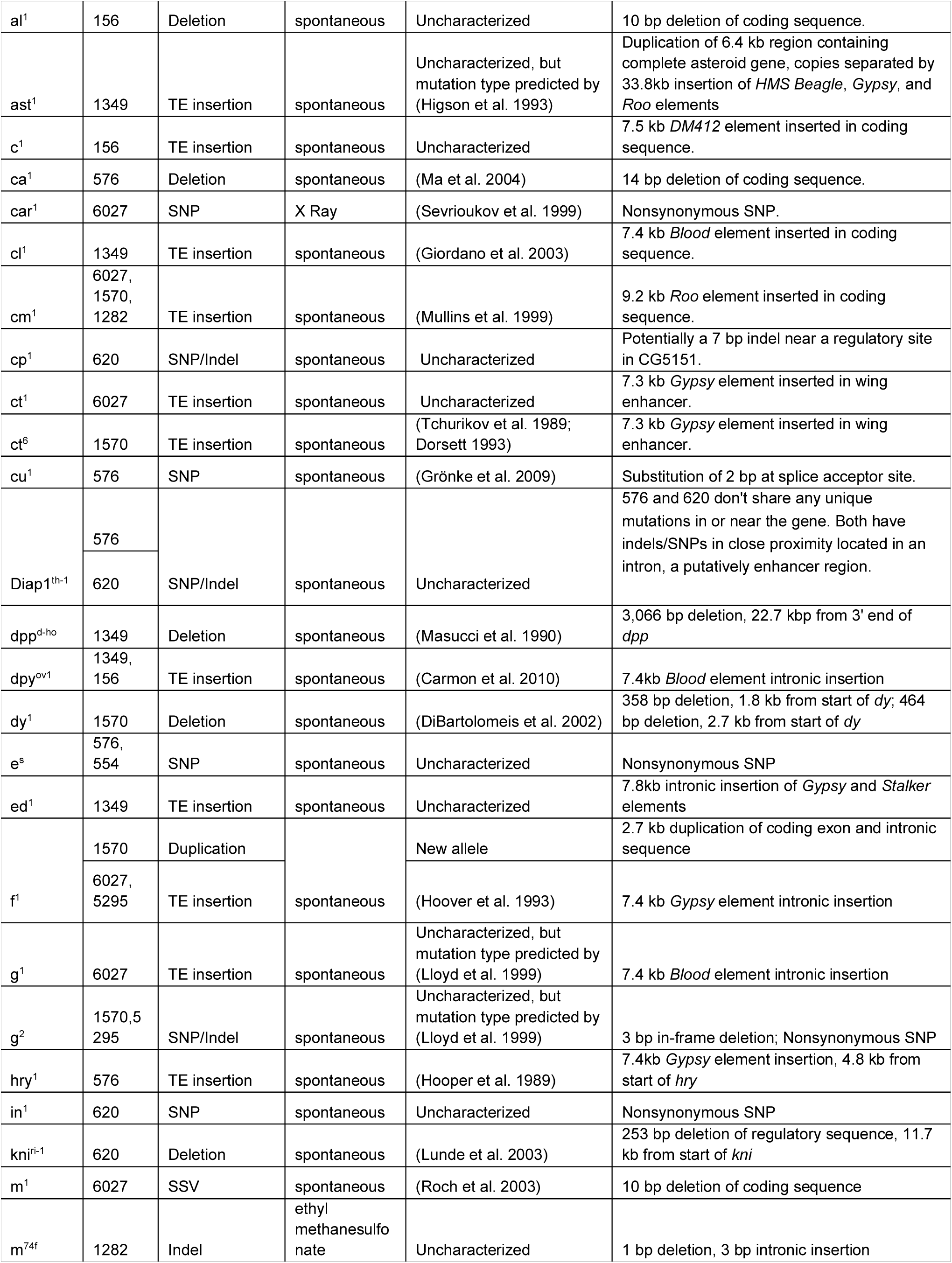
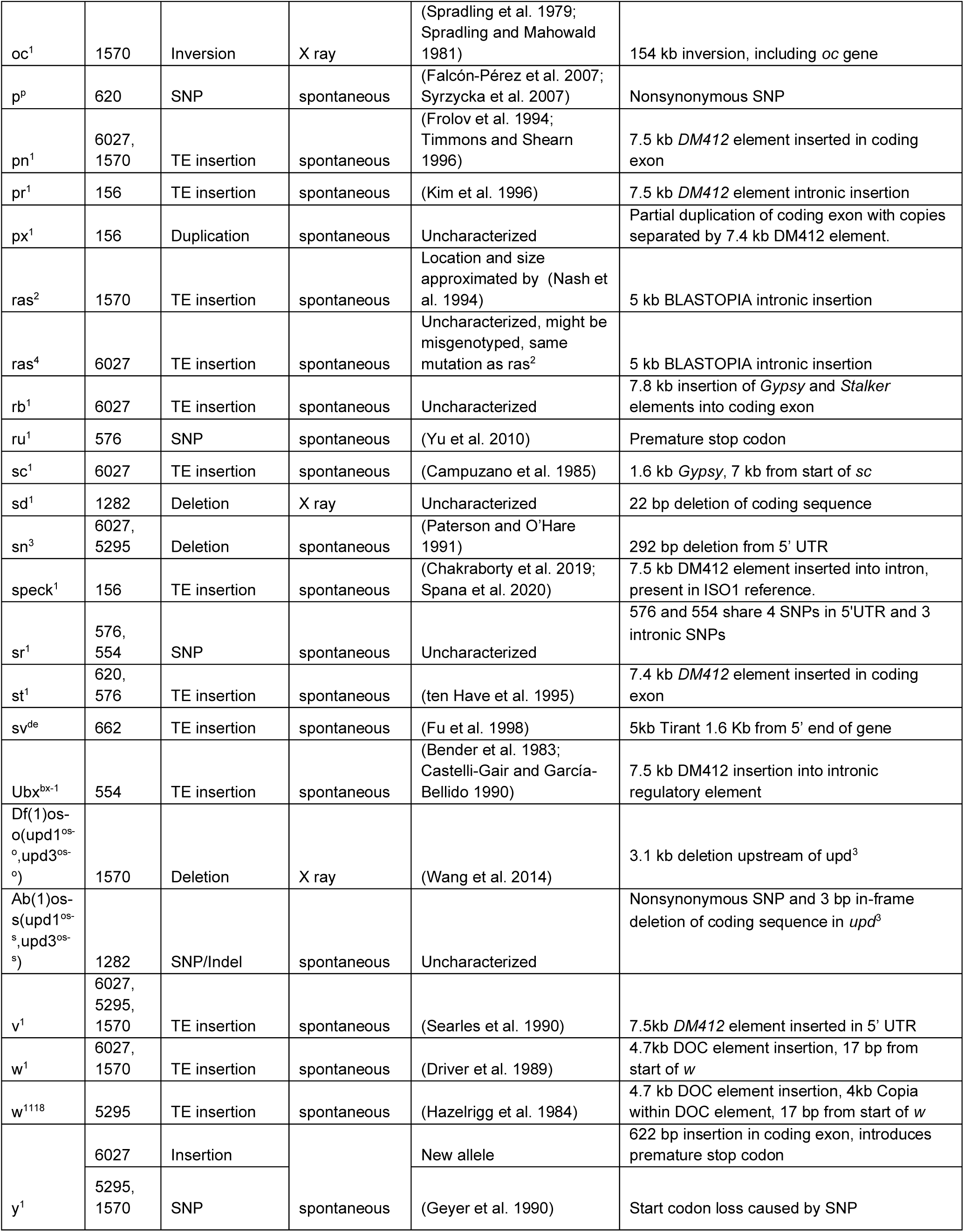
50 Visible Phenotypes and Candidate Mutations.

We identified a candidate for the unmapped *Abnormal leg pattern* (*Ablp*) gene, which bears a mutation associated with the *Dp(2;4)ey^D^* translocation duplication, *Ablp^eyD^*. This gene was previously mapped to a 90 kb region on chromosome 2L, at the source of the sequence translocated onto the 4th chromosome in the *Dp(2;4)ey^D^* mutation. Within this mapped location, strain 662, which carries *Ablp*^eyD^, has an 8 kb *Roo* element insertion into a predicted transcription factor binding site (TFBS) for *paired* (*prd*) (MacArthur et al. 2009) (Fig. 2a), a key regulator of segmental patterning in *Drosophila* development (Kilchherr et al. 1986). This TFBS is located between *drumstick* (*drm*) and *sister of odd and bowl* (*sob*), both members of the odd-skipped gene family involved in leg joint formation (Hao et al. 2003). Using CRISPR-Cas9, we initially generated a full deletion of the predicted regulatory site with two gRNAs, but flies carrying this deletion showed no detectable leg joint phenotype relative to the unedited controls (Supplementary Fig. 5). In contrast, an independent edit that removed 11 bp, designated *prdBS*^Δ1^ for *‘paired* binding site knockout 1’, produced abnormal development of the first tarsal joint - the same segment affected in *Ablp*^eyD^ mutants (Fig. 2a, b) and amplicon sequencing confirmed the precise 11 bp deletion at the target (Supplementary Fig 6). Together, these results support a regulatory role for the sequence adjacent to the *Roo* element insertion site, and suggest that it may interfere with *paired* binding, potentially leading to misregulation of *drm* and or *sob*, causing the tarsal joint abnormalities observed in *Ablp*^eyD^ mutants.

**Figure 2.**
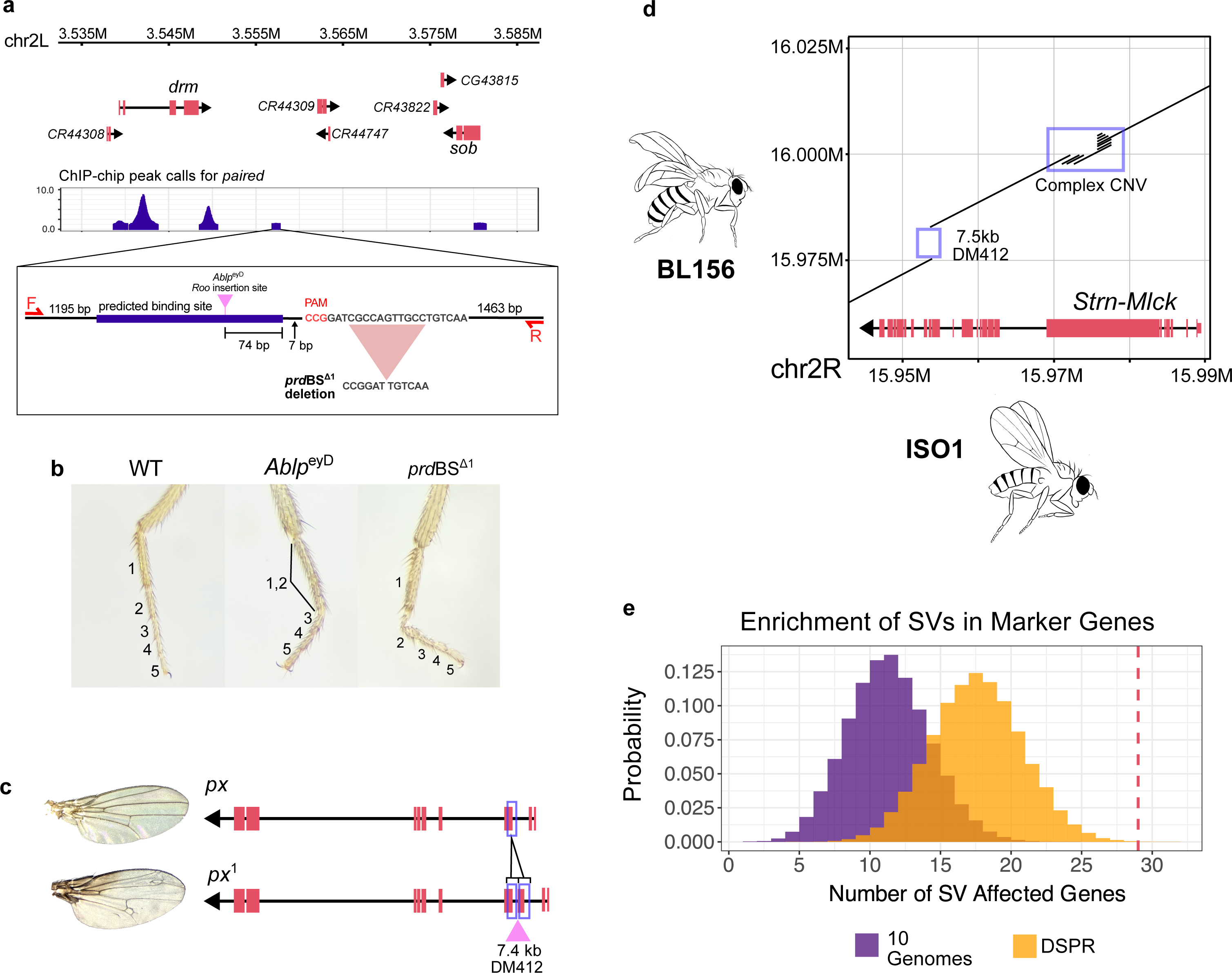
a. Gene model and ChIP-chip binding peaks for the transcription factor *paired*, showing the location of the CRISPR target sequence relative to the predicted binding site (purple) and the 8 kb Roo element insertion present in the *Ablp*^eyD^ strain (pink). *prd*BS^Δ1^ is an 11 bp deletion located 14 bp from the 3’ end of the predicted binding site. Forward and reverse primers (F+R) were designed to amplify the genomic region for genotyping. b. Tarsal segment phenotypes of a wild-type, *Ablp*^eyD^, and homozygous *prd*BS^Δ1^ mutant. The *Ablp*^eyD^ phenotype involves defects at the joint between tarsal segments 1 and 2, typically with partial fusion of the tarsal segments. *prd*BS^Δ1^ shows a defect at the same joint. c. Structure of the *plexus* gene in the ISO1 wild-type allele (top) and strain 156, which carries the *p*x^1^ mutation (bottom). The *p*x^1^ allele has a 1.5 kb partial duplication of an exon, with a DM412 TE insertion between the copies. d. Dot plot alignment between the genomes of strain 156, which carries the curved-wing *c*^1^ mutation, and the ISO1 reference at the gene *Strn-Mlck.* While the complex CNV is found in several strains lacking the c^1^ phenotype, the DM412 insertion is unique to strain 156. e. Monte Carlo distribution of the number of genes bearing an SV in the 10 genomes SV map (purple) and in the DSPR SV map (orange) in samples of 43 genes. In the 43 marker genes associated with the 50 phenotypes analyzed in this study, 29 contain a candidate SV (red line).

Another mutation, px^1^, is associated with increased wing veins at the wing margins and tips. However, the mutation responsible for this gene is yet to be characterized. We found a 1.5 kb tandem duplication that copied the 5’ half of the third exon of *Plexus* (*px*), containing the splice site and intronic sequence, into the second intron. The duplicated segments are separated by a 7.4 kb *DM412* retrotransposon (Fig. 2c). The TE disrupts the reading frame of the duplicated exon and introduces a premature stop codon, likely resulting in a truncated *plexus* protein. Loss of functional *plexus* may impair repression of wing vein development, thereby explaining the ectopic vein phenotype observed in *px*^1^ mutant wings.

The *curved* (c^1^) mutation, associated with the curved wing phenotype, has yet to be mapped to the genome sequence (Supplementary Fig. 8). Previous studies based on deficiency maps indicated 10 genes as potential candidates for *c*, with *Strn-Mlck* being the most probable one (Kahsai and Cook 2018). We identified a 7.5 kb DM412 retrotransposon and a complex duplication, both of which disrupt the coding sequence of *Strn-Mlck*, in strain 156 (Fig. 2d). The TE insertion is absent in the other strains that lack the c^1^ phenotype. Other genes in the interval where c^1^ is mapped did not have any obvious disrupting mutations, suggesting that *Strn-Mlck* is *c*.

A clipped wing phenotype, *clipped* (*cp*), was mapped to a 3-Mbp region on chromosome 3L (see Methods). We further narrowed down the region by crossing flies carrying the *cp*^1^ mutation with deficiency lines (see Methods), reducing the region to a 55 kb window containing 8 protein-coding genes and 3 lncRNAs (Supplementary Fig. 9). One of these genes, CG5151, has been shown to play an important role in wing development. Knockdown of the gene in the posterior imaginal wing disc results in wing notching similar to that observed in *cp*^1^ mutants (Bageritz et al. 2019). Strain 620 shows the clipped phenotype and carries a 7 bp indel at a predicted TF binding site in CG5151. Thus, 6 phenotypes among the 15 without a candidate mutation were associated with SVs, and we infer the rest are caused by SNPs or small indels (Table 2).

### Prevalence of SVs in deleterious phenotypes

We found that 66% (33/50) of the markers are associated with LSVs, and 6% (3/50) are associated with SSVs (Table 2). The remaining phenotypes are associated with SNPs or small indels that disrupt protein-coding sequences or regulatory sequences. Of the phenotypes caused by spontaneous or natural mutations in our dataset, 71% (32/45) are associated with SVs. Similar to previous observations that TEs cause many visible phenotypic changes in *D. melanogaster* (Sankaranarayanan 1988), 46% (23/50) of mutations are associated with TEs. We also find duplication copy number variation (CNV), indels, and inversions underlying the mutations (Table 2).

To determine whether SVs are disproportionately associated with visible phenotypic changes, we compared the prevalence of SVs in the 43 genes linked to the 50 phenotypic markers to their genome-wide distribution in the 10 genome assemblies in which these mutations were identified. We performed a Monte Carlo simulation, randomly drawing 100,000 gene sets matched for gene length to our marker gene set, to generate a null distribution for the expected number of SVs (Methods). Based on this null model, we would expect SVs in 12 genes by chance; however, we observed SVs in 29 of the 43 marker genes, representing an enrichment of 141.67% (p-value = 9.99 × 10^-6^) (Fig. 2e). Because the analyzed genomes were generated by crossing multiple strains to combine marker mutations, their SV content may differ from the spectra of mutations in chromosomes segregating in natural populations. To address this, we repeated the analysis using SV calls from the genome assemblies of 14 inbred strains collected from various geographical locations worldwide (Chakraborty et al. 2019). Based on the abundance of SVs in this population sample, we expect SVs in 18 genes by chance. The observation of 29 genes with SVs in the marker set thus represents a 61.1% enrichment (p-value = 7.90 × 10^-4^).

All 50 phenotypes examined in this study are associated with visible phenotypic changes, and 44 among these have deleterious effects on health and behavior (Supplementary Table 6). For instance, the vermilion eye color mutation(*v*^1^) causes slow and irregular heart rates (Beasley and Dowse 2016); *white* eye color mutations (*w*^1118^ and *w*^1^) are linked to defects affecting mobility, lifespan, and courtship behavior (Krstic et al. 2013; Xiao et al. 2017; Arimoto et al. 2020); Bristle mutants (*forked*) exhibit a reduced response to courtship sounds (Cosetti et al. 2008), and yellow body color mutant (*yellow*) males have lower mating success due to reduced melanization of their sex combs (Massey et al. 2019). Among the 44 markers associated with deleterious fitness effects, 75% (33/44) are associated with an SV.

### Allelic diversity underlying phenotypic changes

Similar phenotypes can often result from distinct mutations in the same gene (Schmidt et al. 2010; King et al. 2014; Chakraborty et al. 2019; GTEx Consortium 2020), though the prevalence of multiple alleles at loci underlying variation in deleterious organismal phenotypes remains unclear. To examine the diversity of molecular alleles underlying the deleterious phenotypes in our dataset, we inspected the genes underlying the 12 phenotypes present in more than one strain. We found multiple alleles involving SVs linked to four phenotypes shared between strains, although three SV alleles among these were previously unknown.

Notably, two phenotypes include classic, well-characterized mutations, such as *white* and *yellow*, which result in white eyes and a yellow body color, respectively (Table 2). Our analysis revealed previously uncharacterized molecular diversity underlying these phenotypes. For example, strains 1570, 5295, and 6027 possess the *y*^1^ mutation, although only 1570 and 5295 have the previously characterized start codon loss mutation caused by a SNP in the initiation codon of the *y* gene (Geyer et al. 1990). The strain 6027 instead has a 622 bp insertion in a protein-coding exon of *y* (Fig. 3a,b). This insertion disrupts the ORF and introduces a premature stop codon, likely leading to a truncated *y* protein. Similarly, *f*^1^ is thought to be caused by an intronic *Gypsy* TE insertion (Hoover et al. 1993). While strains 1570, 5295, and 6027 show the forked mutant phenotype, only strains 5295 and 6027 have the TE. The *f* gene in 1570 has a tandem duplication that copies exonic and intronic sequences and is likely to disrupt the ORF of the gene and produce a mutant phenotype (Fig. 3c, d).

**Figure 3.**
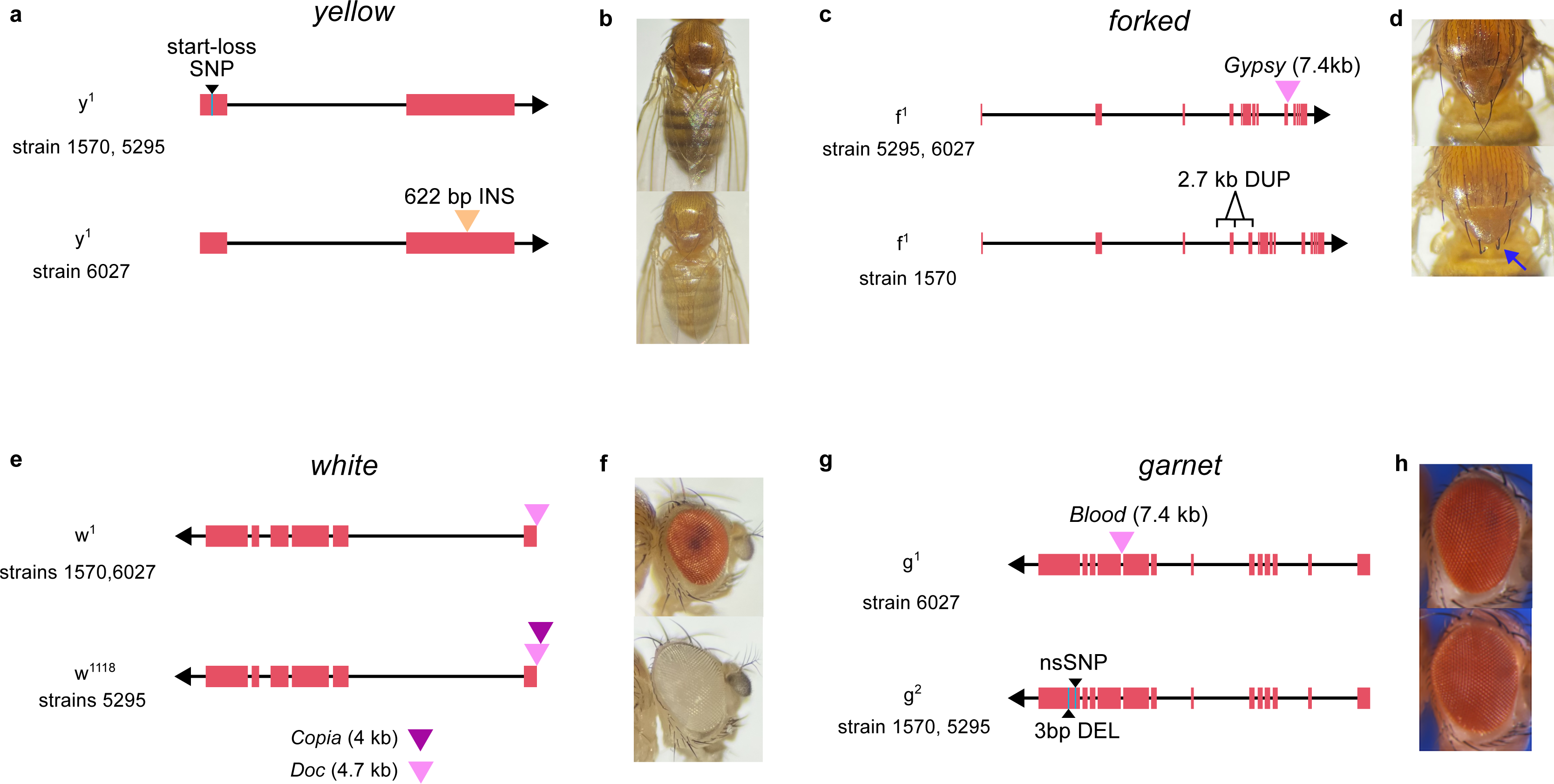
a. Two strains showing the yellow phenotype have the previously characterized SNP which results in loss of the start codon. Strain 6027 instead has an insertion of 622 bp into the second exon which disrupts the reading frame and results in a premature stop. b. Phenotype images of the wild-type (top) and *y*^1^ (bottom) body color. c. The *f*^1^ allele was previously linked to a TE insertion, however, we identify one strain, 1570, lacking the TE which instead has a 2.7kb duplication of a complete exon. d. Phenotype images of wild-type (top) and forked bristles of an *f*^1^ mutant (bottom, blue arrow). e. Two known alleles of the *white* gene are linked to TE insertions near the transcription start site of the gene. Both alleles involve a *Doc* element insertion at the same site, but the *w*^1118^ has a *Copia* element within the *Doc* sequence. f. Phenotype images of wild-type (top) and *w*^1118^ mutant (bottom) eye color. g. Two alleles of the gene *garnet* involve an intronic *Blood* element insertion (*g*^1^) and a nonsynonymous SNP and in-frame deletion in the last coding exon (*g*^1^). h. Phenotype images of the wild-type (top) and *g*^1^ eye color, obtained from FlyBase https://flybase.org/reports/FBrf0220532.html.

We also uncovered the new molecular basis of mutations that are documented as alleles of the same gene. For instance, *w*^1^ and *w*^1118^ are alleles of the *white* gene, both of which are associated with loss of eye pigmentation. The *w*^1^ allele was previously linked to a TE insertion near the transcription start site, and we confirm this by finding a 4.7 kb *Doc* insertion in strains 1570 and 6027 (Figure 3f). While *w*^1118^ was thought to involve a deletion (Hazelrigg et al. 1984), we instead find that strain 5295 carries a *Doc* insertion at the same position as the *w*^1^ allele, with a 3.5 kb Copia element inserted within the *Doc* element (Fig. 3e, f). These insertions occur at the same genomic site, suggesting that recurrent TE insertions at this locus may give rise to the *white* eye phenotype. We also find evidence that distinct mutation types can have similar phenotypic effects. *g*^1^ and *g*^2^ are alleles of the eye color gene *garnet* (*g*). We show that *g*^1^ is associated with an intronic 7.4 kb *Blood* insertion, whereas *g*^2^ is linked to an in-frame three-bp deletion and an nsSNP in the last exon (Fig. 3g, h).

Additionally, two sets of mutations listed as distinct alleles in the stock genotypes may, in fact, share the same molecular basis. *ct*^1^ and *ct*^6^ are both alleles of the *cut* gene, and are associated with wing notches. The molecular basis of *ct*^6^ has been characterized and is caused by a *Gypsy* TE inserted between the *cut* promoter and a distant wing-margin enhancer (Dorsett 1993; Cai and Levine 1997). We found that strains 6027 and 1570, which are genotyped as *ct*^1^ and *ct*^6^, respectively, both carry this same *Gypsy* TE insertion at the same position, with no other unique disruptive mutations in the *cut* gene. Likewise, *ras*^2^ and *ras*^4^ are alleles of the eye color gene *raspberry*. *Ras2* is linked to a 5 kb *Blastopia* insertion, and both strains 1570 and 6027 carry this same TE insertion at the same site, again with no other clear disruptions in the gene. These findings suggest that the recorded genotypes for these stocks may be incorrect or that the alleles may share the same underlying mutation. The presence of phenotypic differences despite identical disruptions could also mean that additional genetic modifiers, located outside of the previously mapped gene regions, may influence the observed phenotypes.

## Discussion

SVs underlie deleterious and adaptive phenotypic changes and have been hypothesized to account for a portion of the missing causal variants in complex trait variation (Manolio et al. 2009; Eichler et al. 2010). However, their overall impact on phenotypic variation, including traits affecting fitness, remains unknown. In *D. melanogaster,* ∼80% of spontaneous visible mutations affecting 12 phenotypes have been shown to be linked to TEs (Green 1988; Sankaranarayanan 1988). However, these phenotypes were already known to involve TEs. Therefore, a systematic investigation of the role of SVs, both TE and non-TE, in an unbiased, defined set of traits has not yet been conducted, leaving their contribution to phenotypic variation unclear. To address this gap, we constructed a comprehensive variant map of 11 strains carrying 50 classic *D. melanogaster* phenotypic mutations widely used as genetic markers. We show that SVs are associated with a disproportionately higher number of phenotypic changes compared to SNPs, including eight cases involving previously undetected SVs. The identification of novel candidate SVs mirrors growing evidence from model organisms (Chakraborty et al. 2019), crops (Chia et al. 2012; Alonge et al. 2020), livestock (Li et al. 2024; Yang et al. 2024), and humans (Abel et al. 2020), indicating that hidden SVs may underlie a wide range of traits, from physiology to behavior. Although the variants underlying these visible phenotypes were not sampled from natural populations, their nature - often single, large effect mutations - is similar to that of phenotypic differences observed in diverse biological contexts. Such mutations underlie well-characterized cases of morphological and behavioral divergences between closely related species or populations (ex. beach mouse color pattern (Hoekstra et al. 2006), stickleback skeletal changes (Shapiro et al. 2004; Colosimo et al. 2005), butterfly mimicry (Naisbit et al. 2003; Kunte et al. 2014)) as well as large effect alleles contributing to crop and livestock domestication and improvement (Andersson 2013; Wills et al. 2013). Our dataset consists of mostly spontaneous mutations identified in lab stocks, however, they can serve as useful models for understanding how large mutations contribute to phenotypic variation.

SVs, particularly TEs and duplicates, show enrichment among low-frequency variants, suggesting stronger purifying selection (Emerson et al. 2008; Cridland et al. 2013; Chakraborty et al. 2019; Abel et al. 2020; Collins and Talkowski 2025; Samano et al. 2025). However, the biological basis for this pattern remains poorly understood. While TE insertions are generally considered disruptive, we do not know what proportion of phenotypic changes affecting fitness are due to TEs or other SVs. Based on published data, we inferred that 44 phenotypes in our study have deleterious fitness effects, with the majority (75%) of these phenotypes being associated with SVs (Supplementary Table 6). Deleterious recessive mutations like these marker phenotypes often segregate at low frequencies in natural populations. Since the allele frequency distribution of SVs in the strains studied here resembles the SFS of SVs in strains from natural populations (Chakraborty et al. 2019), the enrichment of SVs among deleterious phenotypes could offer a biological perspective for population genetic inferences that SVs often exert stronger harmful effects than nonsynonymous SNPs. Additionally, our data shows that similar deleterious phenotypic changes can result from different SV alleles, with the proportion of phenotypes (3/12) exhibiting allelic heterogeneity being similar to the proportion of *D. melanogaster* genes showing SV allelic heterogeneity. Yet, these SVs may evade detection, even when they occur in widely studied genes such as *yellow* and *white*. The prominent role of hidden and multiallelic SVs in deleterious phenotypes underscores their potential significance in disease genetics. As demonstrated by the *D. melanogaster* traits examined here, SVs account for a substantial proportion of harmful alleles and thus may contribute disproportionately to genetic disorders and the unexplained heritability of complex diseases (Manolio et al. 2009; Eichler et al. 2010). The enrichment of SVs among deleterious large-effect phenotypic changes is also consistent with the sizable contribution of SVs towards inbreeding depression and extinction risks, particularly in small populations experiencing weak natural selection (Rogers and Slatkin 2017).

SVs can influence gene structure and function through diverse mechanisms. While duplications of complete genes have long been recognized as drivers of phenotypic variation and adaptations (Hughes 1994; Chakraborty and Fry 2015; Cardoso-Moreira et al. 2016), genomic rearrangements involving partial genes can also have functional and evolutionary consequences. For example, the *jingwei* gene in *Drosophila* originated through the retrotransposition of *Adh* exons into another gene, resulting in a novel exon structure and a new gene function (Long and Langley 1993). Duplications of one or more exons in the dystrophin gene can disrupt protein function and lead to muscular dystrophy (White et al. 2006). Consistent with these examples, we show that partial gene duplications, such as the exonic duplications observed in *plexus* and *forked* mutants, can have functional consequences that shape organismal phenotypes. Furthermore, 27 out of the 50 candidate mutations are located in non-coding regions such as UTRs, introns, and intergenic regions. This observation underscores the role of noncoding variation in shaping phenotypes and is consistent with mapping studies identifying trait-associated loci in non-coding regions (Maurano et al. 2012; Alsheikh et al. 2022; Schipper and Posthuma 2022).

While reference based methods are widely used for variant detection, they frequently miss complex or large SVs, especially in repetitive regions. In addition, mapping short reads to reference genomes often fails to resolve complex SVs involving repetitive sequences such as tandem duplications and TEs (Chakraborty et al. 2018). In contrast, strain-specific assemblies and pangenome approaches offer a more comprehensive view of genetic variation facilitating the discovery of novel SVs (Hickey et al. 2023). For example, the mutation *asteroid*, which causes a rough eye phenotype, involves a complete duplication of the gene followed by insertion of TEs. Although earlier work had predicted the location of this mutation (Higson et al. 1993), our *de novo* genome assembly captures the size and structure of this mutation. Similarly, *px*^1^ is caused by a tandem duplication with a 7.4 kb retrotransposon inserted between the two copies. Such mutations cannot be detected by methods that rely on mapping the orientation of paired-end short reads (Chakraborty et al. 2018). The copia and Doc elements responsible for the *w*^1118^ mutation also exemplify such complexities, where the proximity of the two TE insertions likely thwarted their detection. These examples show assembly- and pangenome-based approaches can reveal functionally important SVs, including novel molecular alleles in highly studied genes, that would be hidden with read mapping alone, misleading the mutational basis of phenotypic variation.

Beyond variant discovery, *de novo* genome assemblies of individual strains facilitate genome annotation, particularly for regions that remain unmapped. *D. melanogaster*, despite being one of the best-annotated metazoan genomes, still contains genes that have not been mapped to the genome sequence (Dean et al. 2022). We examined three such cases—*Ablp*, *clipped*, and *curved—*and identified genes and a regulatory element potentially responsible for the associated mutant phenotypes. These results highlight how long-read assemblies can help resolve persistent gaps in genome annotation and improve the genome-phenotype map, even in such a well-studied model organism as *D. melanogaster.* Thus, our approach of using long reads to link genotypes to phenotypes provides a model for both scientists and educators to discover and annotate functional genetic elements in a laboratory or a classroom, respectively. Inspired by A.H. Sturtevant’s pioneering undergraduate work in genetic mapping, we integrated this model in a resource called Genomics and Long Reads Education (GALORE) that embodies the same spirit of discovery in the modern classroom, while also supporting more advanced training and research applications, such as recombination landscape inference (see Data Availability).

The discovery of the Bar mutation in *Drosophila* provided the earliest evidence supporting the role of genome structural changes in phenotypic variation (Hurles et al. 2008). Since then, SVs have been implicated in several Mendelian and complex diseases as well as adaptations (Merker et al. 2018; Quan et al. 2021; Collins and Talkowski 2025). However, the extent to which SVs contribute to phenotypic variation remains unclear, partly due to the challenges of detecting comprehensive SVs. Comparative genomics using highly contiguous genome assemblies has largely solved that problem, although our understanding of the contribution of SVs in phenotypic variation remains incomplete. Similar to previous findings of enrichment of SVs in candidate genes in QTL mapping experiments, our results suggest a disproportionate role of SVs in large, deleterious changes in phenotypes with both Mendelian and complex genetic basis (Chakraborty et al. 2019). Thus, our results further show that SVs can act as rare alleles of large effects and may account for undetected causal mutations for variation in Mendelian and complex traits, particularly those affecting organismal fitness.

## Materials and Methods

### Fly stocks and DNA extraction

We obtained the *D. melanogaster* stocks from the Bloomington Stock Center (ordered on 12-10- 2023, received on 12-18-2023). The presence of visual markers listed on the stock center website was verified for each strain. We collected 150 females from each stock and extracted high-molecular-weight DNA using the method described by (Chakraborty et al. 2016). Briefly, we flash-froze the flies in liquid nitrogen and ground them into fine powder using a mortar and pestle. We extracted DNA from the fly powder using the Qiagen Blood and Cell Culture Midi Kit and then spooled the DNA at the final stage using a glass hook.

### Library preparation and sequencing

We prepared the ONT library for each strain following the manufacturer’s ligation kit protocol. The initial concentration and total volume of DNA for each sample are provided in Supplementary Table 2. For high molecular weight DNA from strains 2969, 5295, 576, and 1349, we used the PacBio Short Read Eliminator XL kit to remove DNA fragments below 40kb DNA lengths below 40kb. DNA was end-repaired using the NEBNext Companion Module for ONT Ligation Sequencing Kit (New England Biolabs), followed by adaptor ligation with the ONT Duplex-Enabled Ligation Sequencing Kit V14. Libraries were sequenced on R10.4.1 flow cells using a MinION Mk1B for 72 hours.

### Base-calling and assembly

We performed base calling using a Dorado duplex base-calling model on a laptop computer with 64 GB of memory and 2 TB SSD drives. Although each run produced a small proportion of duplex reads, we did not assemble the duplex reads separately due to their low coverage (Supp. Table 3). Raw ONT reads were filtered using Chopper (De Coster and Rademakers 2023), keeping only reads with an average Phred quality score greater than 10 and lengths greater than 10 kbp. We generated a draft assembly of each genome using Hifiasm v0.25.0 (Cheng et al. 2021). Microbial contigs in the draft assembly were identified using Blobtools v1.1 (Laetsch and Blaxter 2017). For input to the Blobtools analysis, we generated a taxonomic annotation file by aligning contigs to the NCBI nucleotide database (downloaded 4-23-2024) using the BLASTn algorithm. The read-mapping input was generated by mapping the quality-filtered ONT reads to the draft assembly using minimap2 v2.26 (Li 2017). Only contigs classified as “Arthropoda” or “no-hit” were retained. Cleaned, draft assemblies were polished using Medaka (https://github.com/nanoporetech/medaka). Polished contigs were scaffolded using *mscaffolder* (Chakraborty et al. 2018) with the release 6.49 of the ISO1 genome assembly (Hoskins et al. 2015) as the reference.

Strain 662 carries the *eyD* mutation, which involves a translocation duplication of a segment from chromosome 2L onto the 4th chromosome. Hifiasm was unable to resolve this complex structural variant, so we used Flye v2.9.3 to generate and inspect the repeat graph (Kolmogorov et al. 2019). The repeat graph was visualized using Bandage (Wick et al. 2015), which indicated that the duplicated sequence may have been collapsed into a single copy (Supplementary Fig. 4a). To recover the entire structure, we manually expanded the collapsed region by exporting the path as a FASTA. We confirmed the new breakpoints by mapping reads back to the duplicated sequence (Supplementary Fig. 4b).

### Assembly quality assessment

To evaluate assembly quality, we first identified potential large-scale misassemblies by remapping long reads to the contig assemblies using minimap2 v2.26 (Li 2017) and visually inspecting regions with abnormal coverage profiles. Assembly contiguity was quantified using QUAST v5.0.2 (Gurevich et al. 2013), based on standard summary statistics (contig N50, L50, contig number, total assembly size). Completeness was assessed using BUSCO v5.7.1 (Simão et al. 2015) with the Diptera ortholog database downloaded 7-8-24.

### Repeat annotation and SV calling

We annotated repeats in each assembly using RepeatMasker v4.1.2 (Smit et al. 2013). To identify SVs, we combined whole-genome alignment and read-mapping approaches. We utilized the Minigraph-Cactus pipeline (Hickey et al., 2023) to construct a pangenome graph, encompassing all forms of genetic variation across the 11 genomes. From the ISO1-based VCF file, we identified mutations located within annotated genes that were unique to strains carrying marker mutations. Additionally, each assembly was aligned to the ISO1 reference genome, specifically the major chromosome arms and the dot chromosomes (X, 2L, 2R, 3L, 3R, 4), using MUMmer v4 (Marçais et al. 2018). Structural differences between assemblies were then classified as insertions, deletions, duplications, or inversions using SVMU (Chakraborty et al. 2019). Finally, we mapped the ONT reads to the ISO1 reference using minimap2 v2.26 (Li 2017) and inspected them in IGV (Thorvaldsdóttir et al. 2013) to confirm that the read data supported SVs detected by whole-genome alignment-based methods. Strains 1570 and 6027 carry an X chromosome balancer, so the assembled X chromosomes are highly fragmented. To identify mutations linked to X chromosome marker genes in these strains we used Sniffles v2.6.1 (Smolka et al. 2024) to identify SVs from reads mapped to the ISO1 reference. Candidate SVs were similarly verified in IGV.

### SNP and indel calling

We used the PEPPER-Margin-DeepVariant pipeline (Shafin et al. 2021), to accurately call SNPs and small indels from the ONT reads mapped to the ISO1 reference. Individual strain VCF files were normalized and merged with BCFtools v1.19 (Danecek et al. 2021), and variant effects were predicted using SnpEff v5.2 (Cingolani et al. 2012).

### CRISPR genome editing

To examine the functional significance of the predicted regulatory region for the unmapped *Ablp* gene, we selected two guide RNAs (gRNA1: 5’-TGATTGCGAAGAAACCTCTG-3’, gRNA2: 5′- TTGACAGGCAACTGGCGATC-3′) flanking the predicted enhancer site using CRISPOR (Concordet and Haeussler 2018) . gRNA2 is positioned 84 bp downstream of the Roo insertion site and near the 3’ end of the predicted *paired* binding site; gRNA1 targets the opposite flank, 806 bp from the Roo insertion site, to enable full-site deletion. As the gRNA2 sequence is located very close to the 3’ end of the predicted *Paired* binding site, deletions resulting from CRISPR- Cas9 mediated DNA breaks may overlap and disrupt the regulatory site. The synthesized gRNAs (Synthego) were incubated with the Cas9 enzyme (Synthego) to form RNP and injected into embryos of the BDSC strain 54591 (Port et al. 2014). The presence of the gRNA sequence was verified by aligning the putative regulatory sequence from the ISO1 reference sequence to the genome sequence of strain 54591 available at https://github.com/chakrabortymlab/DLPD. Embryo injections were performed by GenetiVision Corporation (Stafford, TX). Surviving females were individually crossed to single males and allowed to mate for five days. Following observation of larval activity, females (G0) were genotyped using PCR followed by amplicon sequencing to assess CRISPR-mediated deletions.

To genotype the individuals for CRISPR-mediated deletion alleles, we isolated genomic DNA from single flies using the Monarch Genomic DNA Purification Kit (New England Biolabs) following the manufacturer’s Genomic DNA Extraction from Insects protocol. The genomic DNA was amplified using Q5 polymerase with primers 5′-TCAGCGAGTACAACTCAGCA-3′ and 5′- TTGTTGTCGCTGGAGATTCGA-3′. The PCR amplicons were purified using the Monarch PCR & DNA Cleanup Kit and sequenced with Oxford Nanopore (Plasmidsaurus). The sequencing reads were mapped to the 54591 assembly using minimap2 v2.26 (Li 2017). The read alignments in BAM format were viewed in IGV to examine the CRISPR-induced deletions. We recovered two deletion alleles: a full deletion between the two gRNAs and an 11 bp deletion at the site of the second gRNA. Presence of only reads carrying a single deletion allele was considered as evidence for homozygosity of the deletion, whereas presence of different edited alleles or a mix of edited and unedited alleles was considered as heterozygous.

F1 males and females from G0 females homozygous for the deletion were crossed and genotyped. Due to the unknown genotype of the G0 male, not all F1 males and females were homozygous for the deletion. Thus, we isolated F2 males and females homozygous for the deletion and inspected their leg phenotypes.

### Deletion mapping of *clipped* gene

Strain 620 carries a wing mutation, *cp*^1^, a mutant allele of the *clipped* gene, which has not yet been localized to a precise location in the genome. According to unpublished data on FlyBase (https://flybase.org/reports/FBrf0198635), *clipped* was predicted to reside within a 3.12 Mb region between the genes *asf1* and *st*. To refine the location of the *clipped* gene, we performed deletion mapping using 14 lines, each carrying a chromosomal deletion spanning different portions of the predicted interval (Supplementary Table 5). These deletion stocks were crossed to strain 620 as well as to strain 466, which also carries the *cp*^1^ allele, and the F1s were examined for the wing notching phenotype associated with *cp*^1^.

Eleven deletions complemented the *cp*^1^ phenotype, indicating that the mutation lies outside the regions deleted in those lines. Three deletions failed to complement, suggesting that these *overlapping* deletions uncover the *cp*^1^ mutation. From the overlap of the three non-complementing deletions, we defined a minimal candidate interval of 54,992 bp on chromosome 2L (Supplementary Fig. 7b). Within this interval, we searched for mutations unique to strain 620 as candidate mutations responsible for the *cp*^1^ phenotype.

### Enrichment of SVs in mapped genes

To determine whether the number of marker genes with candidate SVs was significantly higher than the genome-wide distribution of SVs, we generated a null distribution using our map of SVs in 10 of the genomes. We excluded SVs from strain 2969 from this analysis, as we did not include the *Bar*^1^ mutation among our 50 markers due to its known origin as an SV. We considered SVs larger than 100 bp and identified euchromatic protein-coding genes located within 1,000 bp of any SV. To construct a null distribution, we randomly sampled 43 genes (the number of genes linked to the 50 phenotypes) from the genome, controlling for gene length by matching the distribution of gene lengths in the marker gene set to those in the sampled sets using a kernel density estimation (KDE) and rejection sampling approach. This sampling procedure was repeated 100,000 times. The *p*-value was calculated using the following formula, p=(r+1)/(n+1) (North et al. 2002), where n is the total number of replicates and r is the number of replicates in which the number of genes affected by SVs is equal to or larger than the observed number of marker genes with candidate SVs. This analysis was repeated with a map of euchromatic SVs in the Drosophila Synthetic Population Resource (DSPR), a panel of 14 isogenic lines derived from globally diverse populations (Chakraborty et al. 2019).

## Data availability

All genome assemblies have been deposited to NCBI (Bioproject accession PRJNA1214913). All reads are deposited in the NCBI Short Read Archive (SRA) (Supp. Table 3). All scripts for genome assembly and analysis are available at https://github.com/chakrabortymlab/biol450-2024. Additional step-by-step instructions to carry out the genomic analysis reported here is also available at (https://github.com/chakrabortymlab/GALORE).

## Acknowledgments

We thank all students of the undergraduate Genomics course (BIOL 450) in the Department of Biology at Texas A&M University who took the course in the spring 2024 semester for their assistance with data collection and analysis. We also thank Anthony Long and Trevor Millar for helpful suggestions and feedback on the manuscript. We are grateful to Kevin Cook for providing us with an initial list of BDSC stocks with important visual markers. We also thank High-Performance Research Computing at Texas A&M University for providing the computational resources used in this study. This work was supported by funding from the Department of Biology at Texas A&M University, Texas A&M University startup grant, and the National Institutes of Health grant (R00GM129411) to M.C.

