## Supplementary Materials for "Structural variants are enriched in deleterious visible phenotypes in *Drosophila*"

### Supplementary Figures

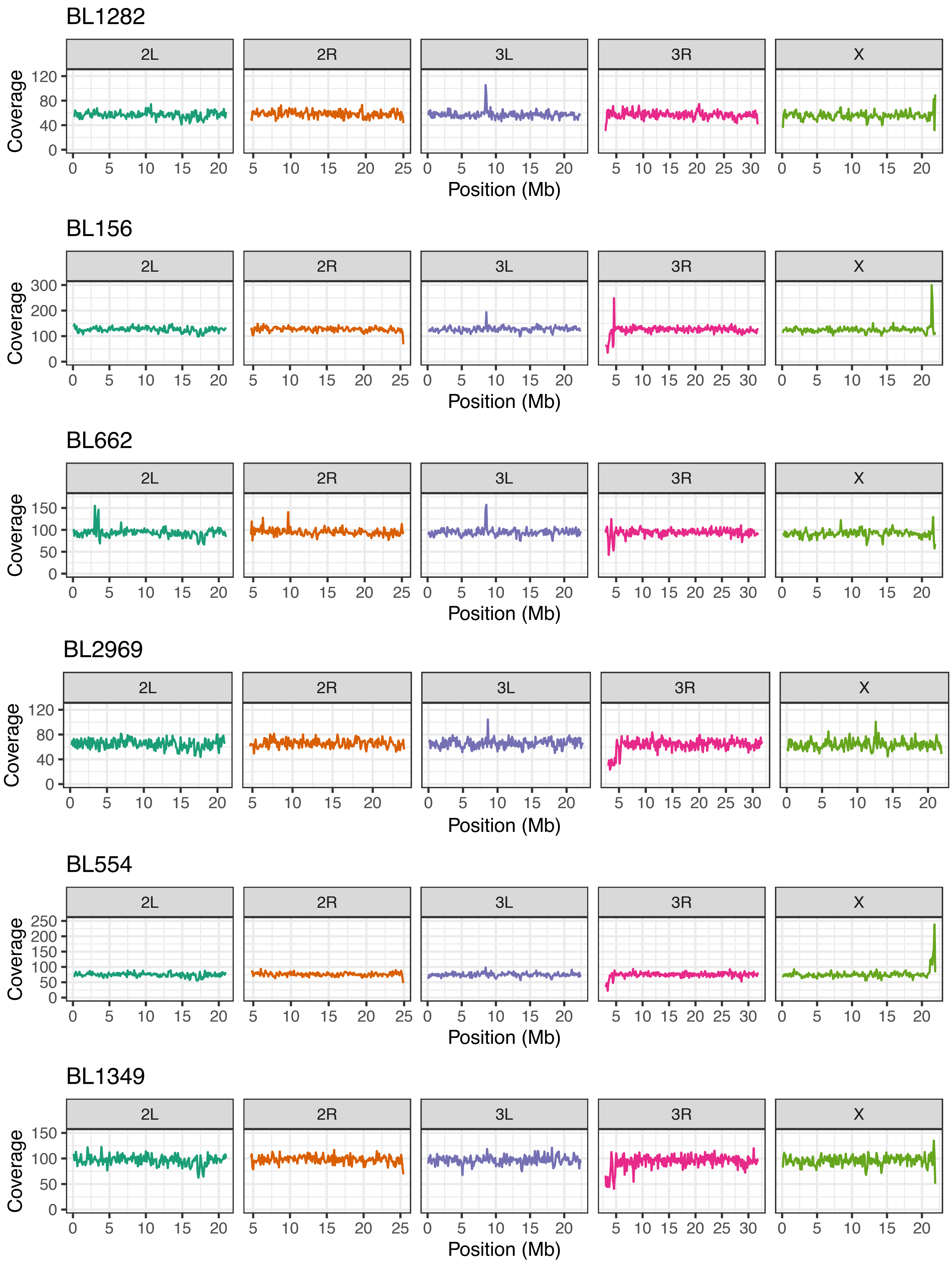

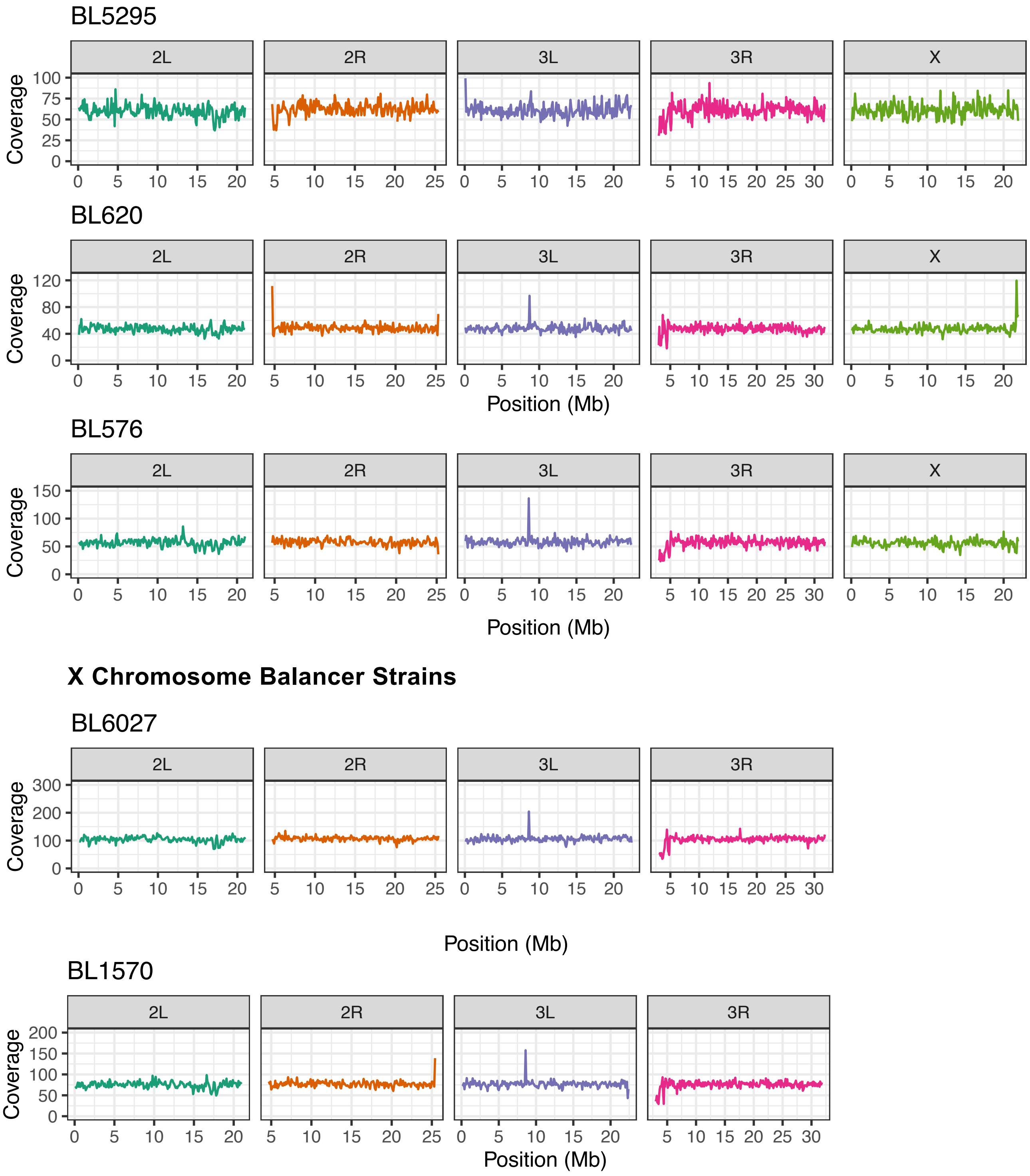


**Supplementary Figure 1.** Read-depth profiles after aligning raw reads to each strain's scaffolded assembly show near-uniform coverage across the major chromosome arms, with no extended dips or spikes indicative of collapsed repeats, consistent with the absence of major misassemblies.

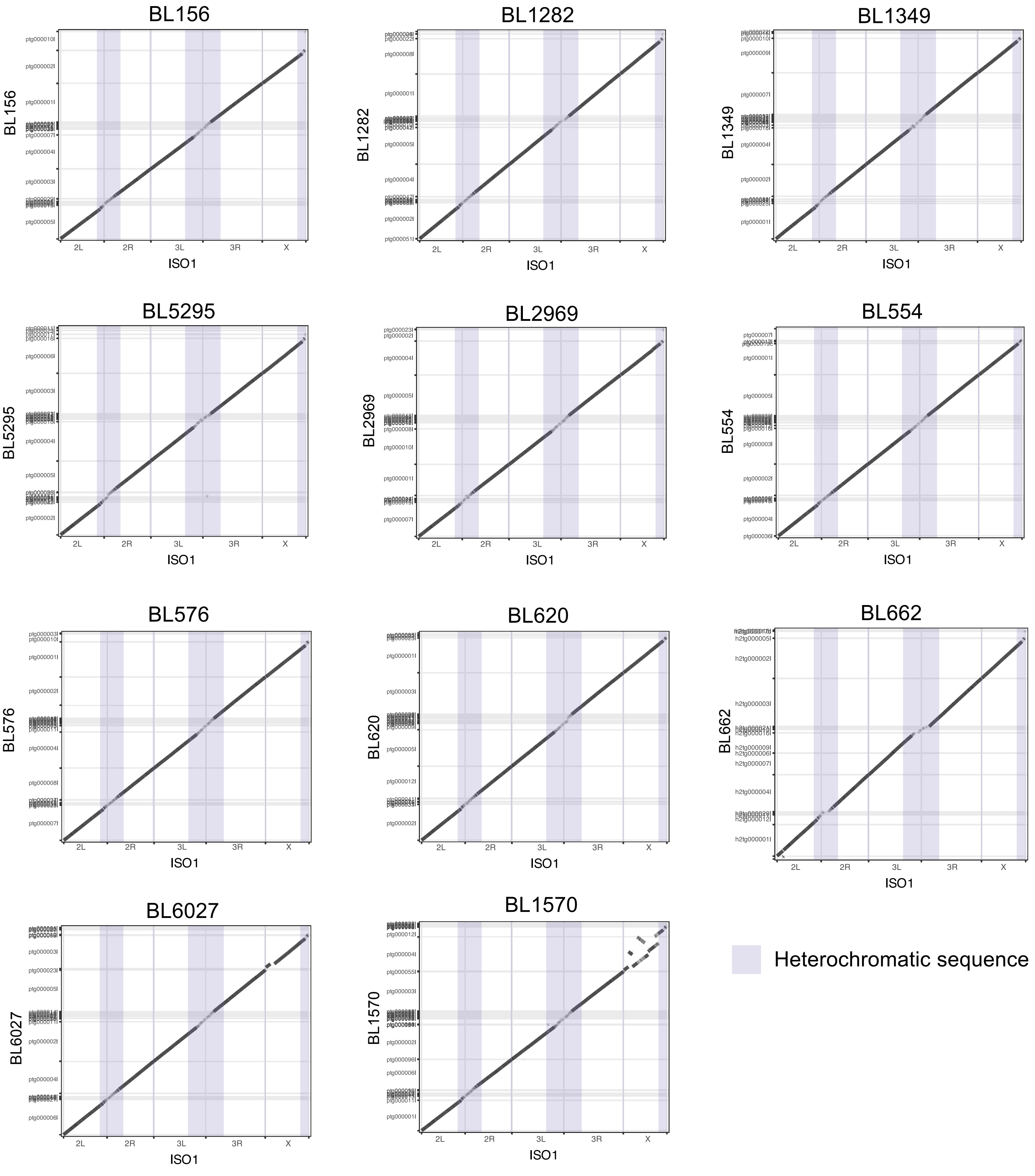


**Supplementary Figure 2.** Alignment dot plots between the ISO1 reference genome and the 11 *de novo* genome assemblies (unscaffolded contigs). Repeats are masked in both reference and query assemblies to show the overall alignment pattern.

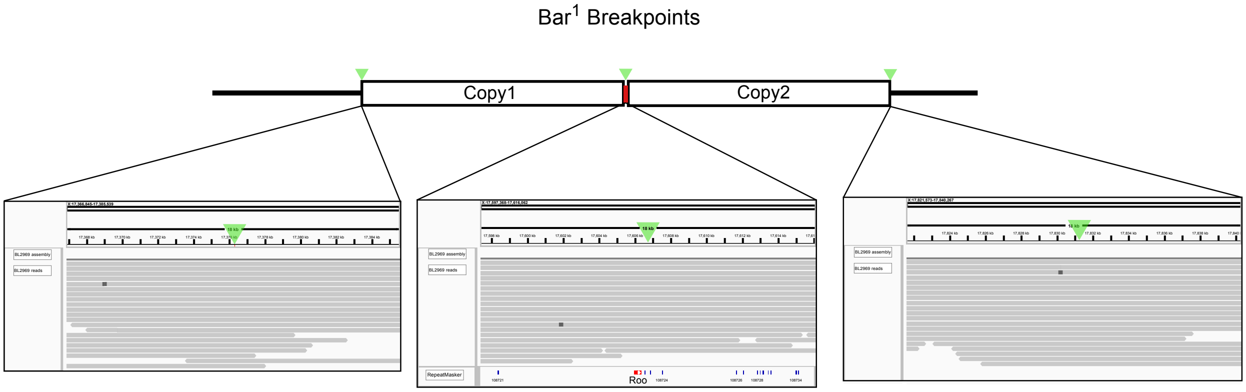


**Supplementary Figure 3.** Read alignment tracks showing read support for the breakpoints of the *Bar*[1] tandem duplication in our assembly for strain 2969. RepeatMasker annotation at the midpoint between the duplicated copies shows the *Roo* element insertion, which is hypothesized to result in unequal crossing over.

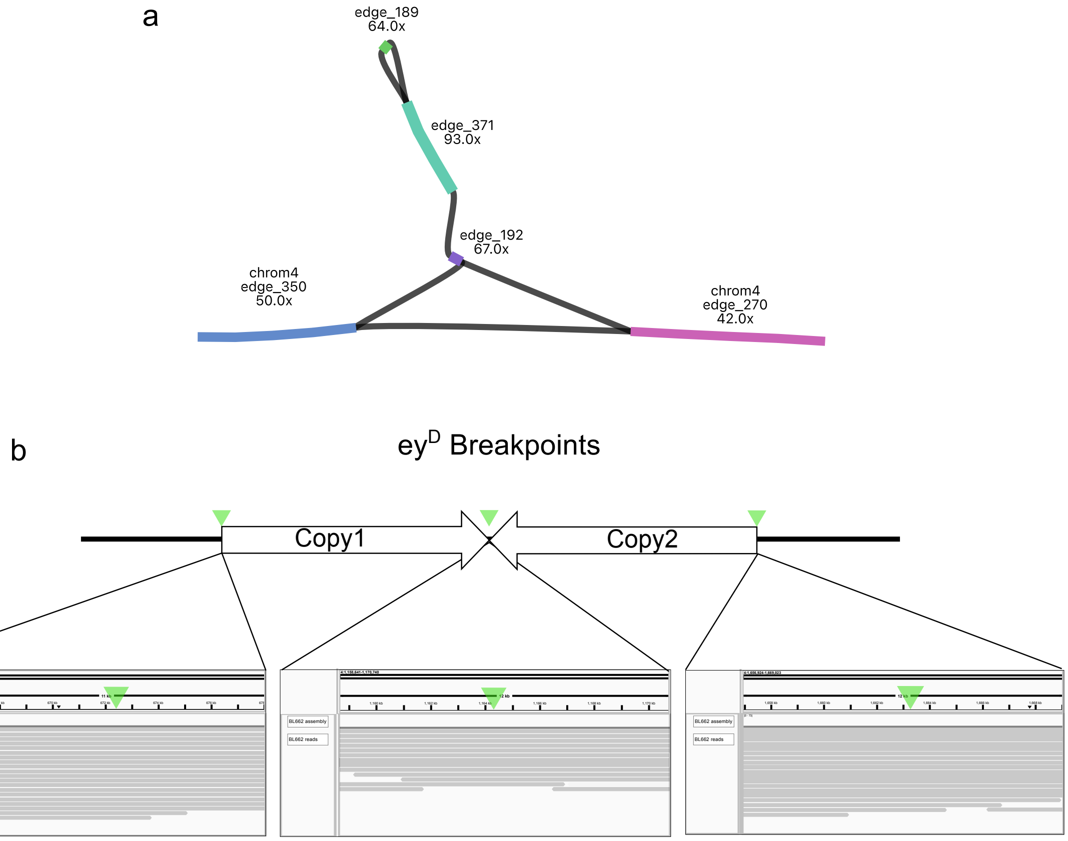


**Supplementary Figure 4.** a. Bandage plot showing the Flye repeat graph at the *ey*[D] translocation-duplication region on chromosome 4 of strain 662. In the assembled genome, the duplication is collapsed, as indicated by the doubled coverage at edge_371. The duplication was manually resolved by exporting the path: edge_350 → edge_192 → edge_371 → edge_189 → edge_371 → edge_192 → edge_270.

b. Breakpoints were validated by confirming read support of the sequence in the corrected genome assembly.

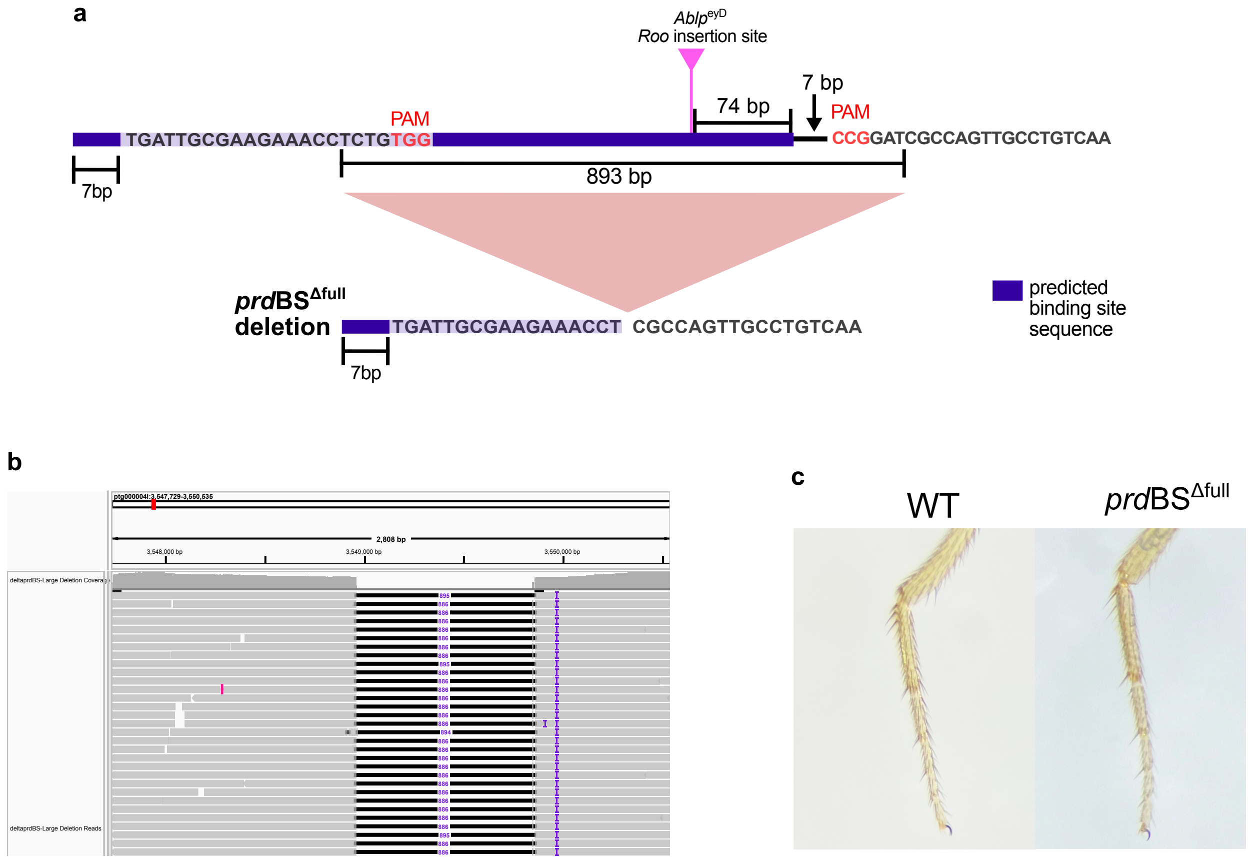


**Supplementary Figure 5.** a. Two guide RNA sequences were designed to produce full site deletion of the predicted binding site for *paired* (purple), in which an 8 kb Roo element (pink) inserts in strain 662, which carries the *Ablp*$^{eyD}$ phenotype. b. Reads from PCR amplification of the targeted locus were aligned to the 54591 genome, showing an 886 bp deletion between the two guide RNA sites. c. Flies carrying the full deletion showed no observable leg joint phenotype compared to the unedited individuals.

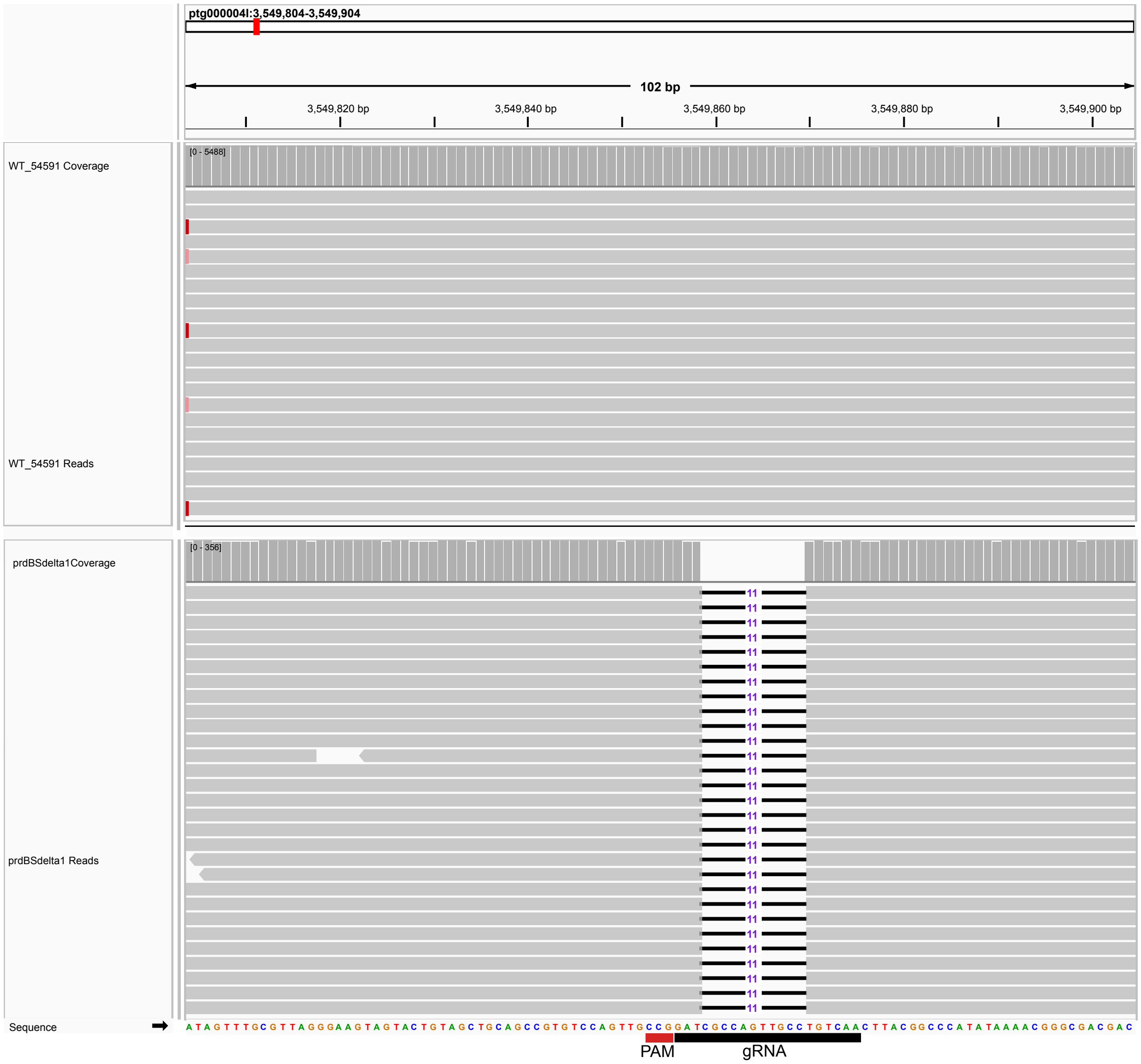


**Supplementary Figure 6.** Reads from PCR amplification of the targeted locus were aligned to the 54591 genome. Top: From a wild-type individual, reads show alignment with no indels across the guide RNA (gRNA) target and PAM, consistent with the 54591 sequence. Bottom: Edited individual displaying the leg-joint phenotype: reads show an 11-bp deletion spanning the predicted Cas9 cut site (3 bp upstream of the PAM), here designated *prd*BS$^{\Delta1}$.

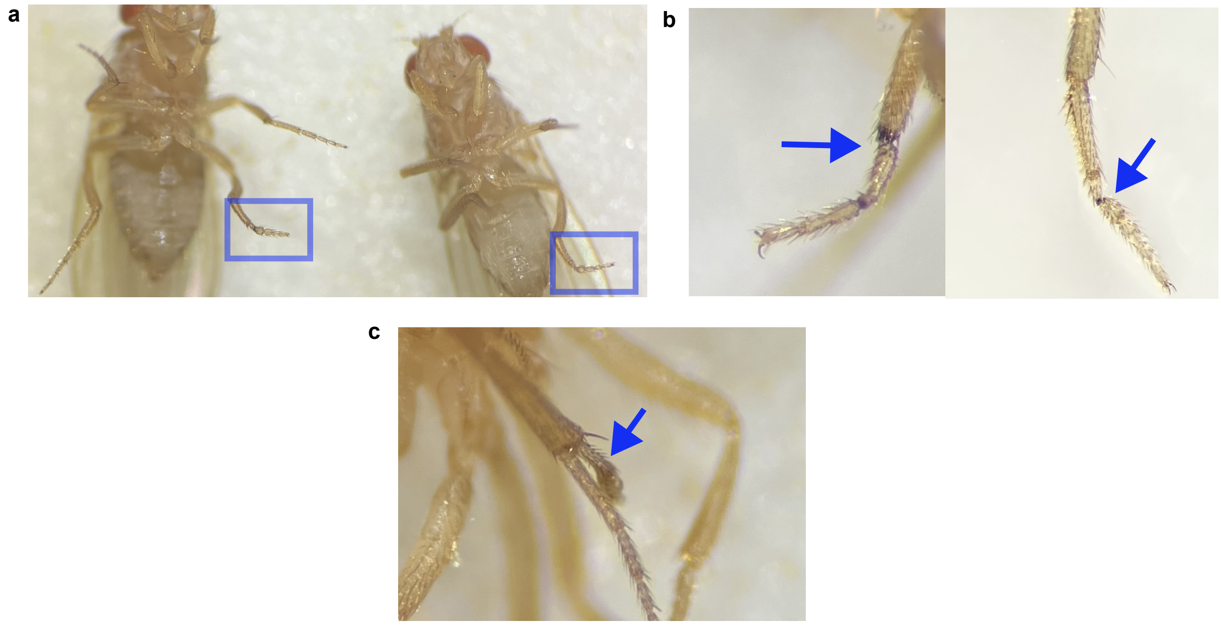


**Supplementary Figure 7.** Phenotype images of transgenic flies carrying CRISPR-mediated deletion at a predicted transcription factor binding site for *paired*. a. Two females displaying the most common phenotype, a pinched tarsal segment. b. Pinched tarsal segments often turned black, indicative of necrotic tissue. c. A single female exhibited a duplicated tarsal segment growing from the tibia.

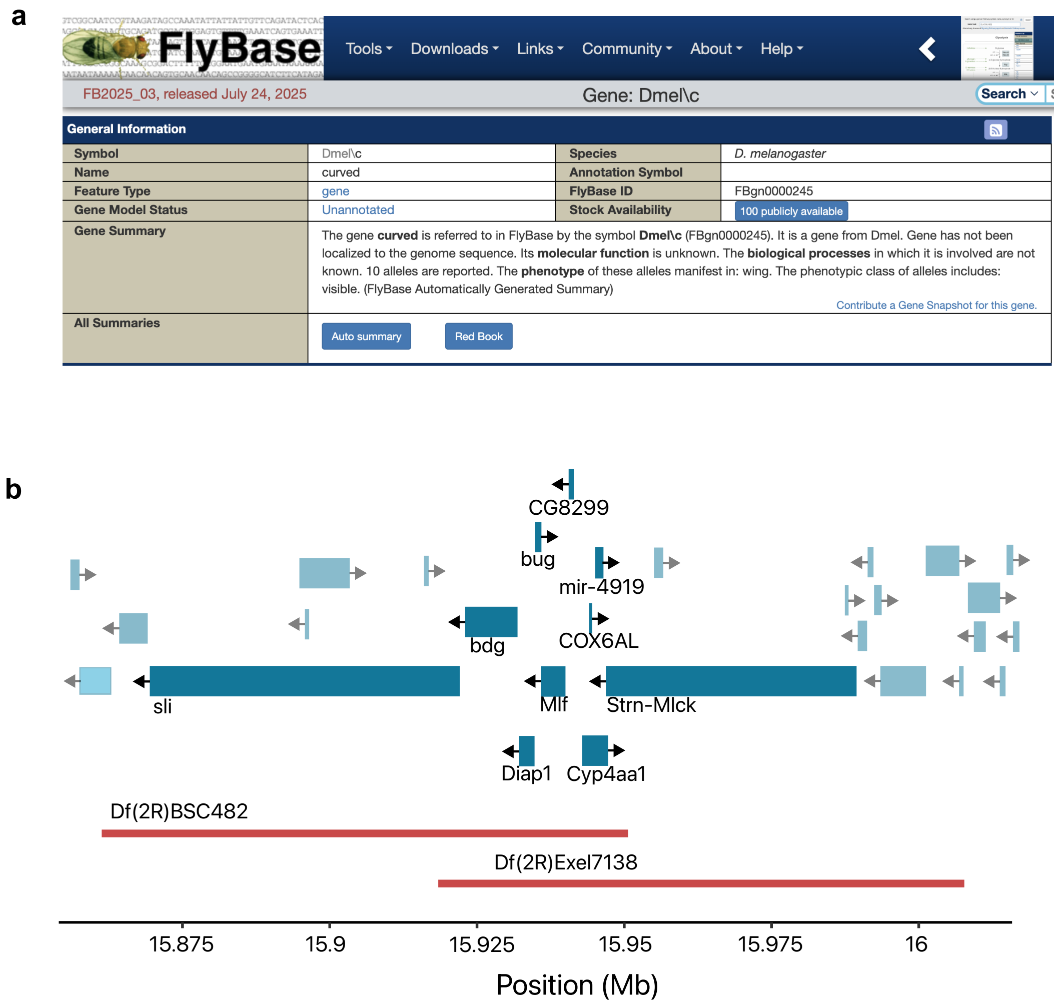


**Supplementary Figure 8.** a. FlyBase page for the gene curved, which is not yet mapped to a genomic location. b. Deletion mapping from a previous study (Kahsai and Cook 2018) narrowed the location of *curved* to a set of 10 genes, with *Strn-Mlck* identified as the most probable candidate.

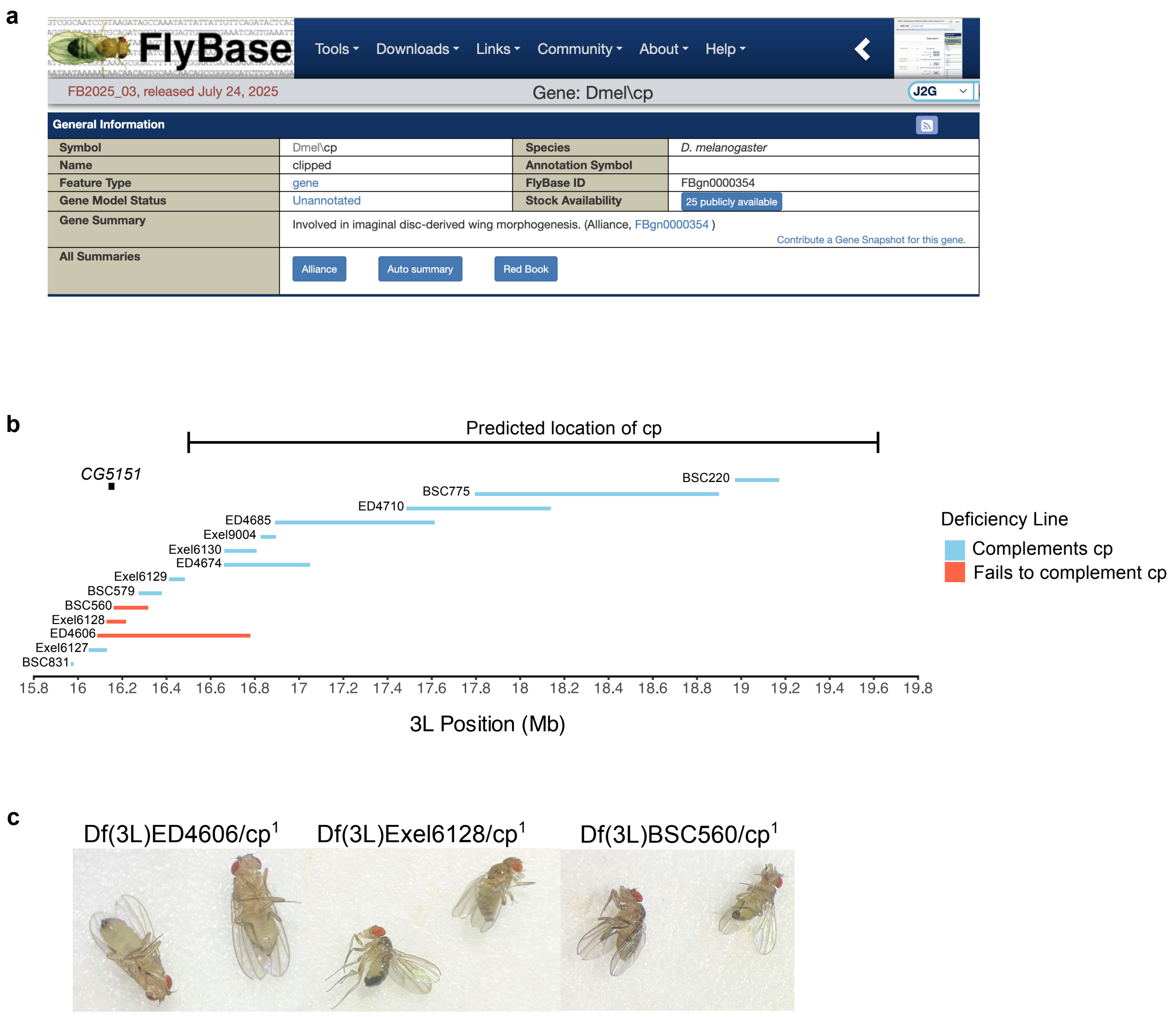


**Supplementary Figure 9**. a. FlyBase page for the gene *clipped*, which is not yet mapped to a genomic location. b. Deletion mapping performed in this study using deficiency lines overlapping or located near the 3.12 Mb region predicted to contain *clipped*. Three deletions failed to complement the *clipped* phenotype, and based on their minimal overlap, the candidate region was narrowed to a 55 kbp interval containing *CG5151*, a gene with a known role in wing development. c. Representative F1 males and females from deficiency line crosses showing failure to complement the $cp^1$ (clipped) phenotype.

### Supplementary Text

#### Additional Information for Previously Uncharacterized Candidate Mutations

*aristaless,* al[1]

Strain 156 has a 10 bp deletion in the last coding exon of *aristaless*, which is the only disruptive mutation that is unique to this strain.

*asteroid,* ast[1]

The breakpoints of this mutation were approximated in a previous publication (Higson et al. 1993) with restriction mapping and cloning. Higson predicted that the mutation was an insertion greater than 21 kb, about 3 kb upstream of the transcription start site of the gene. The genome assembly shows a complete duplication of the *asteroid* gene and a 33.8 kb insertion consisting of *HMS Beagle*, *Gypsy*, and *Roo* elements between the copied region.

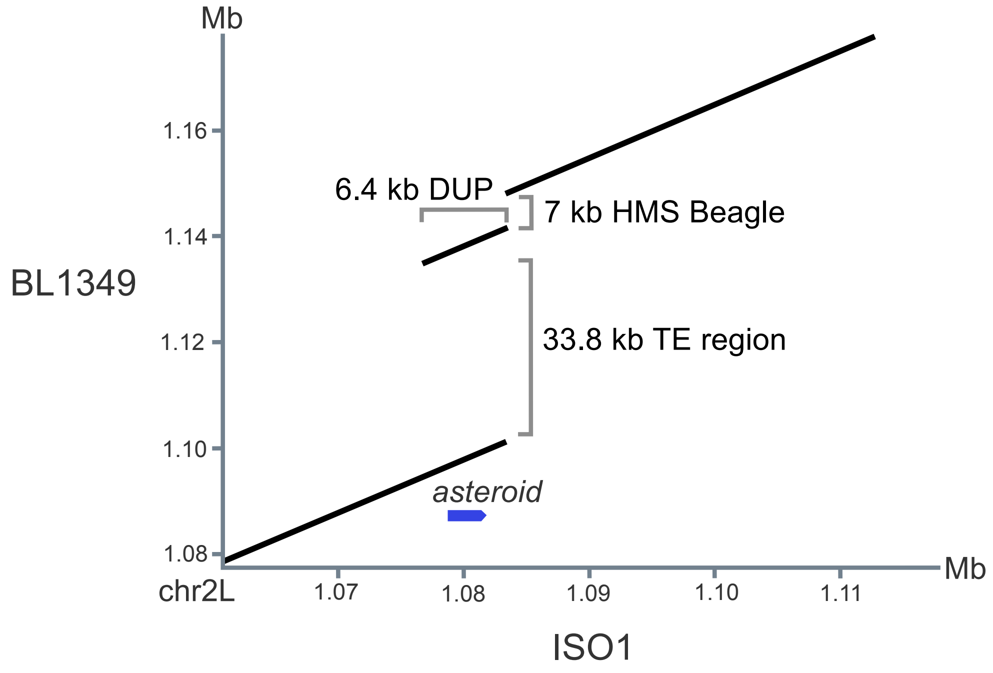


Diap1[th-1] also known as *thread*[1]

Both 620 and 576 show the same *thread* phenotype, where the aristae are lacking lateral branches. Despite this, the strains lack any shared mutations to the gene *Diap1* that are not present in other strains lacking the phenotype. Strain 576 has a nonsynonymous SNP, however, this is not present in 620. Previous study of *thread* mutants postulated that this phenotype might be a regulatory mutation, as it seems to affect only one of the transcript variants, possibly one with a role in aristal antennae development (Cullen and McCall 2004). We do not observe any SVs affecting the *Diap1* gene, so we infer that a SNP or small indel could be the candidate mutation. 620 and 576 both have several intronic SNPs and small indels in close proximity to each other, which affect a predicted regulatory sequence (https://flybase.org/reports/FBsf0000921025).

*ebony, e*[s] or *sooty*

Strains 576 and 554, which both carry the *sooty* allele, have a nonsynonymous SNP in the *ebony* gene which is not present in strains lacking the phenotype.

*echinoid, ed*[1]

Strain 1349 has a 7.8 kb insertion of *Gypsy* and *Stalker* transposable elements, which is not present in other strains lacking the phenotype.

*garnet,* g[1] and g[2]

Southern Blot analysis of g[1] and g[2] mutants (Lloyd et al. 1999), indicated that the g[1] allele is due to a large insertion and g[2] is likely a small point mutation. We find that strain 6027, which carries g[1], has a 7.4 kb Blood element insertion into an intron. Strains 5295 and 1570 are both g[2] and share two coding variations: a 3bp in-frame deletion and nonsynonymous SNP.

in[1]

Strain 620 has a nonsynonymous SNP at 3L: 20,369,329 which is the only disruptive mutation to the *in* gene that is unique to this strain

*miniature* m[74f]

Strain 1282 has several SNPs and indels unique to the strain in the miniature gene, which consist of 3 bp indels in the intron, possibly affecting a regulatory element (https://flybase.org/reports/FBsf0000873514).

*plexus*, px[1]

Strain 156 harbors several large mutations in the *plexus* gene, including a 3.5 kb "deletion" corresponding to the absence of a TE present in the reference genome, a 7.3 kb *Quasimodo* insertion within an intron, a 1.5 kb partial duplication of a coding exon, and a 7.4 kb *DM412* insertion located between the duplicated segments. While the *Quasimodo*, duplication, and *DM412* insertions are unique to strain 156, we predict that the variants disrupting the coding sequence are the likely causative mutations.

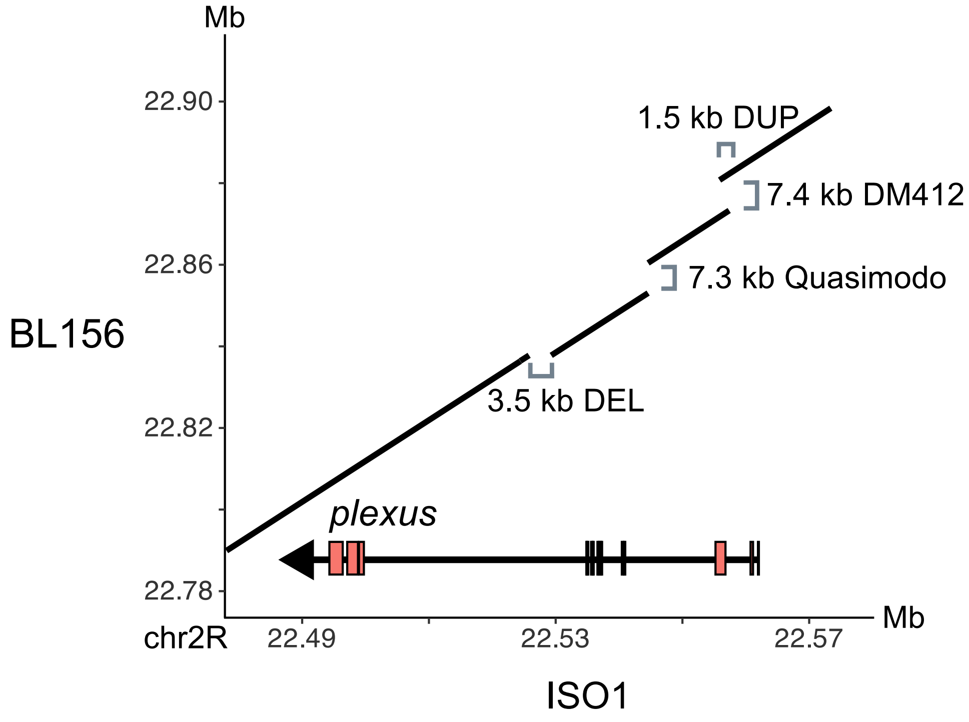


*ruby*, rb[1]

Strain 6027 has a 7.8kb Gypsy1/Stalker2 insertion into a coding exon of *ruby* which is found only in this strain.

sd[1]
Strain 1282 has a 22 bp deletion which removes part of the coding sequence from a single transcript (FBtr0346799), and is unique to this strain.
*stripe,* sr[1]
Strains 576 and 554 share four SNPs in the 5' UTR, two SNPs in an intron, and a 4 bp deletion in an intron, which are not found in other strains lacking the *stripe* phenotype. No SVs were identified within the gene or within 1 kb so we inferred that one or more of the SNPs or indels are the causative mutation.
Ab(1)os-s (upd1[os-s],upd3[os-s])
Strain 1282 has three unique, likely disruptive mutations to the gene *upd*[3], including a disruptive in-frame (3bp) deletion, and a nonsynonymous SNP. The gene *upd1* has no obviously disruptive mutations.

### Supplementary Tables

Supplementary Table 1: Bloomington stock numbers of the sequenced strains and their genotype

| Strain (BL#) | Genotypes |
|---|---|
| 156 | al[1] dpy[ov1] Adc[b-1] pr[1] c[1] px[1] speck[1] |
| 554 | p[p] Ubx[bx-1] sr[1] e[s] |
| 576 | ru[1] hry[1] Diap1[th-1] st[1] cu[1] sr[1] e[s] ca[1] |
| 620 | Diap1[th-1] st[1] cp[1] in[1] kni[ri-1] p[p] |
| 662 | sv[de]/Dp(2;4)ey[D], AbIp[eyD]: ey[D] |
| 1282 | Ab(1)os[s], cm[1] m[74f] sd[1] upd1[os-s] upd3[os-s] |
| 1349 | ast[1] dpp[d-ho] ed[1] dpy[ov1] cl[1] |
| 1570 | Df(1)os[o], y[1] pn[1] w[1] cm[1] ct[6] sn[3] oc[1] ras[2] v[1] dy[1] g[2] f[1] upd1[os-o] upd3[os-o]/FM6 |
| 2969 | Bar[1] |
| 5295 | y[1] w[1118] sn[3] v[1] g[2] f[1]/Dp(1;Y)y[+] |
| 6027 | y[1] ac[1] sc[1] pn[1] w[1] rb[1] cm[1] ct[1] sn[3] ras[4] v[1] m[1] g[1] f[1] car[1]/FM6 |

Supplementary Table 2. Sequencing Statistics

| | SRE kit used? | DNA sequenced (ug) | Bases sequenced (Gb) | Read N50 | %duplex | average coverage |
|---|---|---|---|---|---|---|
| BL1570 | | 254.1 | 12.65 | 12614 | 6.11 | 90.39 |
| BL6027 | | 265.44 | 16.39 | 9272 | 6.25 | 117.08 |
| BL662 | | 265.8 | 15.63 | 9,597 | 6.33 | 111.68 |
| BL156 | | 296.16 | 19.82 | 8,566 | 7.11 | 141.6 |
| BL1282 | | 287.92 | 8.86 | 11,520 | 7.04 | 63.3 |
| BL2969 | yes | 262.14 | 10.29 | 32,293 | 8.9 | 73.51 |
| BL554 | | 203.52 | 11.51 | 17,067 | 7.9 | 82.19 |
| BL5295 | yes | 286.3 | 9.89 | 24,636 | 10.05 | 70.66 |
| BL576 | yes | 275.2 | 9.45 | 20,883 | 7.37 | 67.5 |
| BL620 | | 285.48 | 7.74 | 17,712 | 11.85 | 55.26 |
| BL1349 | yes | 245.88 | 15.77 | 22,059 | 9.12 | 112.62 |

Supplementary Table 3. SRA Accessions

| Bloomington Strain ID (BL#) | SRA Accession | Bioproject ID | Biosample Accession |
|---|---|---|---|
| 662 | SRR32117938 | PRJNA1214913 | SAMN46390171 |
| 554 | SRR32117933 | PRJNA1214913 | SAMN46390169 |
| 1349 | SRR32117939 | PRJNA1214913 | SAMN46389945 |
| 6027 | SRR32117931 | PRJNA1214913 | SAMN46390170 |
| 156 | SRR32117936 | PRJNA1214913 | SAMN46389946 |
| 5295 | SRR32117932 | PRJNA1214913 | SAMN46390627 |
| 576 | SRR32117934 | PRJNA1214913 | SAMN46390156 |
| 2969 | SRR32117935 | PRJNA1214913 | SAMN46389952 |
| 620 | SRR32117930 | PRJNA1214913 | SAMN46390618 |

| | | | |
|---|---|---|---|
| 1570 | SRR32117937 | PRJNA1214913 | SAMN46389947 |
| 1282 | SRR32117940 | PRJNA1214913 | SAMN46389944 |

Supplementary Table 4. Assembly statistics

| | Contig Assembly | | | Scaffolded Assembly | | | Diptera BUSCOs |
|---|---|---|---|---|---|---|---|
| | Size (Mbp) | #contigs | Contig N50 (Mb) | #scaffs | Scaffold N50 (Mb) | Ns per 100 kbp | Complete |
| BL1570 | 159.8 | 97 | 21.8 | 31 | 27.1 | 1.43 | 99.4 |
| BL6027 | 151.4 | 69 | 23.9 | 48 | 28.3 | 1.32 | 98.6 |
| BL662 | 135.7 | 22 | 21.8 | 13 | 26.6 | 0.66 | 99.2 |
| BL156 | 154.9 | 41 | 23.8 | 22 | 29.1 | 1.16 | 99.4 |
| BL1282 | 146 | 56 | 24.1 | 33 | 27.4 | 1.51 | 99.4 |
| BL2969 | 151.8 | 55 | 24 | 36 | 29.5 | 1.19 | 99.3 |
| BL554 | 150.4 | 51 | 24.1 | 27 | 28.6 | 1.53 | 99.4 |
| BL5295 | 154.3 | 54 | 24.3 | 31 | 27.1 | 1.43 | 99.4 |
| BL576 | 151.8 | 44 | 21.7 | 21 | 29 | 1.45 | 99.5 |
| BL620 | 151.9 | 51 | 24.3 | 28 | 28.9 | 1.45 | 99.4 |
| BL1349 | 153.4 | 78 | 24.2 | 55 | 29.3 | 1.43 | 99.4 |

Supplementary Table 5. *Clipped* mapping

| BDSC Strain | Deficiency Line | Start | End | Complemented cp? |
|---|---|---|---|---|
| 29021 | Df(3L)BSC831/TM6C | 15,967,776 | 15,979,964 | No |
| 7606 | Df(3L)Exel6127 | 16,046,940 | 16,129,654 | No |
| 8078 | Df(3L)ED4606 | 16,087,484 | 16,780,123 | Yes |
| 7607 | Df(3L)Exel6128 | 16,129,654 | 16,217,328 | Yes |
| 25122 | Df(3L)BSC560/TM6B | 16,162,336 | 16,318,079 | Yes |

| | | | | |
|---|---|---|---|---|
| 25413 | Df(3L)BSC579/TM6C | 16,274,238 | 16,378,604 | No |
| 7608 | Df(3L)Exel6129 | 16,411,693 | 16,483,557 | No |
| 8098 | Df(3L)ED4674 | 16,661,284 | 17,049,418 | No |
| 7609 | Df(3L)Exel6130 | 16,661,291 | 16,806,648 | No |
| 7937 | Df(3L)Exel9004/TM6B | 16,826,256 | 16,895,610 | No |
| 64121 | Df(3L)ED4685 | 16,891,076 | 17,612,170 | No |
| 8100 | Df(3L)ED4710 | 17,487,463 | 18,139,299 | No |
| 27347 | Df(3L)BSC775 | 17,795,144 | 18,898,326 | No |
| 9697 | Df(3L)BSC220 | 18,972,562 | 19,171,268 | No |

Supplementary Table 6. Deleterious effects of mutations

| # | Mutation | Origin | Phenotypic Defects | Defects References |
|---|---|---|---|---|
| 1 | ac[1] | spontaneous | Lack of sensory bristles | DOI:10.1007/BF01681532 |
| 2 | Adc[b-1] | spontaneous | dark color, other cuticular defects and target recognition | DOI:10.1016/j.gene.2005.03.013 |
| 3 | al[1] | spontaneous | claws and arista are reduced in size, females exhibit reduced mating success | DOI: 10.1242/dev.127.20.4315, 10.1016/s0003-3472(71)80025-8, DOI:10.1016/s0003-3472(71)80025-8 |
| 4 | ast[1] | spontaneous | abnormal eye morphology, muscular defects | DOI: 10.1242/dev.120.7.1731, DOI: 10.1242/dev.00843 |
| 5 | c[1] | spontaneous | wings curved | Lindsley, D.L., Grell, E.H. (1968) |
| 6 | ca[1] | spontaneous | reduced mating success in males | DOI: 10.1038/hdy.2011.60 |
| 7 | car[1] | X Ray | reduced lifespan in males | DOI:10.1534/genetics.106.065011 |
| 8 | cl[1] | spontaneous | no other defects observed besides abnormal eye color | |
| 9 | cm[1] | spontaneous | males have reduced lifespan, retinal response to light defective | DOI:10.1534/genetics.106.065011, DOI:10.1371/journal.pbio.1001847 |
| 10 | cp[1] | spontaneous | clipped wing margins, reduced viability | Lindsley, D.L., Grell, E.H. (1968) |
| 11 | ct[1] | spontaneous | defective wing margins and deformed antennae | DOI: 10.1242/dev.124.17.3241 |
| 12 | ct[6] | spontaneous | defective wing margins and abnormal gravitaxis | DOI:10.1093/genetics/160.4.1481, DOI:10.1007/BF01074308 |
| 13 | cu[1] | spontaneous | wings are curved upward, defective | DOI:10.1266/jjg.31.321 |

| | | | | |
|---|---|---|---|---|
| 14 | Diap1[th-1] | spontaneous | aristae missing lateral branching, abnormal gravitaxis | DOI:10.1007/BF01074308,DOI:10.1007/BF01074308,DOI:10.1016/0092-8674(95)90150-7 |
| 15 | dpp[d-ho] | spontaneous | held out wings, defects on the wing blade | DOI:10.1101/gad.4.11.2011, |
| 16 | dpy[ov1] | spontaneous | defective, dumpy wings | DOI:10.1016/j.devcel.2015.06.019 |
| 17 | dy[1] | spontaneous | reduced wing size | FlyBase page FBal0003265 |
| 18 | e[s] | spontaneous | abnormal body color, defects in activity and visual response | DOI:10.3109/0167706910906621 3 |
| 19 | ed[1] | spontaneous | eyes rough and misrotated ommatidia | DOI:10.1534/g3.117.300289, DOI:10.1242/dev.038422 |
| 20 | f[1] | spontaneous | abnormal bristles, reduced response to courtship sounds | DOI:10.1177/00034894081170110 6, |
| 21 | g[1] | spontaneous | reduced lifespan in males, disturbed walking and orientation | DOI:10.1534/genetics.106.065011, DOI:10.1093/genetics/155.1.213 |
| 22 | g[2] | spontaneous | abnormal courtship behavior, possibly affects vision | DOI:10.1093/hmg/ddp555 |
| 23 | hry[1] | spontaneous | ectopic wing bristles, minor defects in segmentation | DOI:10.1093/genetics/111.3.463, DOI: 10.1002/j.1460-2075.1993.tb05854 .x |
| 24 | in[1] | spontaneous | abnormal hair polarity, incomplete extra leg joints | DOI:10.1093/genetics/160.4.1535 |
| 25 | kni[ri-1] | spontaneous | missing wing vein, wings are warped and blunt | DOI:10.1534/genetics.110.118695 , FlyBase Page |
| 26 | m[1] | spontaneous | strong reduction of wing size | DOI:10.7717/peerj.12175, DOI:10.1016/j.devcel.2009.11.009 |
| 27 | m[74f] | ethyl methanesulfonate | abnormally long wings | FlyBase page, Craymer, L. (1980). [New mutants report.] D. I. S. 55(): 197--200. |
| 28 | oc[1] | X ray | lack ocelli and associated bristles, underdeveloped brain, female sterile when homozygous | DOI:10.1093/genetics/139.4.1623, DOI:10.1007/s004270100149, DOI:10.1007/s003590050387 |
| 29 | p[p] | spontaneous | no known physiological effects other than eye color | |
| 30 | pn[1] | spontaneous | defects in nervous system development, increased mortality | DOI:10.3389/fncel.2019.00076, |
| 31 | pr[1] | spontaneous | no known physiological effects other than eye color | |
| 32 | px[1] | spontaneous | extra wing veins, structural defects in wing veins | DOI: 10.1242/dev.126.23.5207 |
| 33 | ras[2] | spontaneous | dark ruby eyes | DOI: 10.1007/BF00285747 |
| 34 | ras[4] | spontaneous | dark ruby eyes, female sterility | DOI:10.1007/BF02191716 |
| 35 | rb[1] | spontaneous | shortened male lifespan, defective eye pigment granules, reduced vision, behavior abnormalities | DOI: 10.1534/genetics.106.065011, DOI: 10.1007/pl00008688 |
| 36 | ru[1] | spontaneous | heart abnormalities, misrotation of ommatidia, rough eyes | DOI:10.1371/journal.pgen.1000969 , DOI:10.1016/j.devcel.2004.09.001, PMID: 10887159 |

| | | | | |
|---|---|---|---|---|
| 37 | sc[1] | spontaneous | loss of bristles on multiple body parts, loss of chemoreceptors on wing margins | DOI: 10.1007/BF00260865, DOI:10.1093/genetics/131.2.353 |
| 38 | sd[1] | X ray | jagged wings, defects in bristles and halteres | DOI:10.1007/s004270050201, DOI:10.7554/eLife.00999, FlyBase page |
| 39 | sn[3] | spontaneous | deformed sensory bristles, abnormal axon morphology (curled) | DOI10.1242/dev.036517, DOI:10.1523/JNEUROSCI.2106-06.2006 |
| 40 | speck[1] | spontaneous | no known defects besides darkening of wing hinge | |
| 41 | sr[1] | spontaneous | flightless, muscular and cuticular defects | DOI:10.1093/genetics/119.1.105, DOI: 10.1073/pnas.92.22.10344 |
| 42 | st[1] | spontaneous | reduced lifespan and reduced locomotor ability | DOI:10.1242/jcs.216697 |
| 43 | sv[de] | spontaneous | sterile, sensory defects, uncoordinated | ISBN:0070-7333, DOI:10.1242/dev.126.10.2261 |
| 44 | Ubx[bx-1] | spontaneous | Halteres are transformed into wings as well as several other limb defects | DOI: 10.1002/j.1460-2075.1986.tb04497.x |
| 45 | Df(1)os-o(upd1[os-o],upd3[os-o]) | X ray | reduced eye size, abnormal haltere position | DOI:10.1016/j.ydbio.2014.09.015 |
| 46 | Ab(1)os-s(upd1[os-s],upd3[os-s]) | spontaneous | no known deleterious effects, improved immune response to bacteria | DOI: 10.1038/ncomms14642 |
| 47 | v[1] | spontaneous | abnormal, slow heart beat | DOI:10.1002/jez.2057 |
| 48 | w[1] | spontaneous | decreased copulation rate | DOI:10.1371/journal.pone.0001391, DOI:10.1038/s41598-017-08155-y |
| 49 | w[1118] | spontaneous | deterioration of climbing ability, reduced courtship, retinal degradation under certain light conditions | DOI:10.3390/ijms222312967, DOI:10.1371/journal.pone.0077904, DOI:10.1091/mbc.E09-10-0917,DOI:10.3390/ijms222312967 |
| 50 | y[1] | spontaneous | reduced mating success in males | DOI:10.7554/eLife.49388, DOI:10.1534/genetics.105.045666 |

Supplementary Table 7. Candidate mutations coordinates (tsv)
